# Prolactin and prolactin receptor expression in the HPG axis and crop during parental care in both sexes of a biparental bird (*Columba livia*)

**DOI:** 10.1101/2021.07.13.452208

**Authors:** Victoria S. Farrar, Rayna M. Harris, Suzanne H. Austin, Brandon M. Nava Ultreras, April M. Booth, Frédéric Angelier, Andrew S. Lang, Tanner Feustel, Candice Lee, Annie Bond, Matthew D. MacManes, Rebecca M. Calisi

## Abstract

During breeding, multiple circulating hormones, including prolactin, facilitate reproductive transitions in species that exhibit parental care. Prolactin underlies parental behaviors and related physiological changes across many vertebrates, including birds and mammals. While circulating prolactin levels often fluctuate across breeding, less is known about how relevant target tissues vary in their prolactin responsiveness via prolactin receptor (*PRLR*) expression. Recent studies have also investigated prolactin (*PRL*) gene expression outside of the pituitary (i.e., extra-pituitary *PRL*), but how *PRL* gene expression varies during parental care in non-pituitary tissue (e.g., hypothalamus, gonads) remains largely unknown. Further, it is unclear if and how tissue-specific *PRL* and *PRLR* vary between the sexes during biparental care. To address this, we measured *PRL* and *PRLR* gene expression in tissues relevant to parental care, the endocrine reproductive hypothalamic-pituitary-gonadal (HPG) axis and the crop (a tissue with a similar function as the mammalian mammary gland), across various reproductive stages in both sexes of a biparental bird, the rock dove (*Columba livia*). We also assessed how these genes responded to changes in offspring presence by adding chicks mid-incubation, simulating an early hatch when prolactin levels were still moderately low. We found that pituitary *PRL* expression showed similar increases as plasma prolactin levels, and detected extra-pituitary *PRL* in the hypothalamus, gonads and crop. Hypothalamic and gonadal *PRLR* expression also changed as birds began incubation. Crop *PRLR* expression correlated with plasma prolactin, peaking when chicks hatched. In response to replacing eggs with a novel chick mid-incubation, hypothalamic and gonadal *PRL* and *PRLR* gene expression differed significantly compared to mid-incubation controls, even when plasma prolactin levels did not differ. We also found sex differences in *PRL* and *PRLR* that suggest gene expression may allow males to compensate for lower levels in prolactin by upregulating *PRLR* in all tissues. Overall, this study advances our understanding of how tissue-specific changes in responsiveness to parental hormones may differ across key reproductive transitions, in response to offspring cues, and between the sexes.

## 1. Introduction

In animals that exhibit offspring care, an array of physiological changes must occur to facilitate parental behaviors. This transition requires synchronized changes at many physiological levels, from the brain (Bridges, 2015; Champagne and Curley, 2012) to the reproductive organs (Stiver and Alonzo, 2009). Hormones facilitate those changes, including those produced by the reproductive or hypothalamic-pituitary-gonadal (HPG) axis, through their pleiotropic effects on multiple behavioral and physiological traits (Ketterson et al., 2009; Zera and Harshman, 2001). Similarly, tissue responsiveness to hormones, via hormone receptor expression, must also change to produce a synchronized parental phenotype across the brain and periphery (Ball and Balthazart, 2008).

One such hormone, prolactin, plays an important role in parental behavior across vertebrates, but is particularly important in birds (Angelier and Chastel, 2009). Best known for promoting lactation in mammals, prolactin also underlies the onset and maintenance of parental behaviors in birds such as incubation onset, offspring defense and provisioning (Angelier and Chastel, 2009; Buntin, 1996; Smiley, 2019). Circulating prolactin is released by the anterior pituitary gland, and acts upon specific receptors to trigger signaling pathways in target cells. Once in circulation, prolactin acts upon its receptor (PRLR) to activate secondary messenger cascades in target cells, such as the signal transducer and activator of transcription 5 (STAT5) pathway (Austin and Word, 2018; Freeman et al., 2000). Prolactin receptors have been identified in nearly every tissue type in both mammalian and avian species, reinforcing its role in multiple physiological and behavioral processes including reproduction, immune function, and homeostasis (Nagano and Kelly, 1999; Zhou et al., 1996). Additionally, evidence for local prolactin expression beyond the pituitary gland (i.e. extra-pituitary prolactin) has been identified in tissues ranging from the gonads to the mammary glands and the brain (Ben-Jonathan et al., 1996; Marano and Ben-Jonathan, 2014).

While circulating prolactin often increases during parenthood, less is known about how concordant responsiveness to prolactin changes in the brain. In female rats, *PRLR* mRNA increases in some hypothalamic nuclei, and hypothalamic responsiveness to prolactin (measured via STAT5 phosphorylation downstream of the PRLR) increases with reproductive experience (Anderson et al., 2006; Sjoeholm et al., 2011). In birds, brain responsiveness to prolactin increases during breeding compared to non-breeding individuals of multiple species (Buntin and Buntin, 2014; Smiley et al., 2021), and prolactin binding varies seasonally, including during breeding (Smiley et al., 2020). However, how hypothalamic responsiveness to prolactin varies during transitions *within* the breeding cycle remains less studied.

Understanding these subtle changes in *PRLR* expression is important, as changing neural responsiveness to prolactin may prepare behavioral and endocrine systems for the onset of offspring care. For instance, in mammals, prolactin and placental lactogen secretion increases during pregnancy to facilitate lactation and maternal adaptations for postnatal care (Bridges, 2015). In birds, prolactin increases after egg laying to promote incubation behavior with a subsequent increase around hatching to facilitate chick brooding and provisioning in species that exhibit these behaviors (Angelier et al., 2016b; Buntin, 1996; Smiley, 2019). Thus, changes in *PRLR* expression with offspring cues and fluctuating plasma prolactin levels may play a role in prolactin’s facilitation of parental behaviors.

Beyond the brain, peripheral endocrine systems can also respond to prolactin and may influence behavior through altered hormone regulation. *PRLR* gene and protein expression has been documented in the pituitary gland, gonads and other tissues across vertebrates (Aoki et al., 2019; Nagano and Kelly, 1999; Zhou et al., 1996). Prolactin can have an “anti-gonadal” effect in some species, where high circulating levels inhibit sex steroid release and gonadal function (Grattan, 2018; Meier, 1969; Moult and Besser, 1981), which may serve to maintain parental efforts on the current brood rather than continuing breeding or starting a new clutch (Angelier et al., 2016b). These effects may be modulated in part by prolactin’s effects on pituitary gonadotroph cells in the release of luteinizing hormone (LH) or follicle-stimulating hormone (FSH), or by direct action on sex steroid production in the gonads (Bachelot and Binart, 2007). Any of these diverse effects on the HPG axis, and ultimately, reproductive behaviors, would depend upon a tissue’s function and ability to respond to prolactin. Thus, measuring how *PRLR* varies across the HPG axis during parental care is key to understanding how prolactin may exert pleiotropic effects during breeding.

Further, local prolactin expression in the brain and other tissues may also vary during parental care and play an autocrine/paracrine role in hormone regulation (Ben-Jonathan et al., 1996; Marano and Ben-Jonathan, 2014). Extra-pituitary prolactin (*ePRL*) gene expression has been measured in various tissues, including the brain, gonads and mammary glands, though its specific role and function remains unclear (Ben-Jonathan et al., 1996; Marano and Ben-Jonathan, 2014). While there is some debate whether *ePRL* becomes a functional protein (Grattan and Le Tissier, 2015), hypophysectomized rats have been shown to have immunoreactive prolactin protein in their brains (DeVito, 1988), giving evidence that bioactive prolactin can be locally translated beyond the pituitary. Characterizing if, and how, *ePRL* expression changes in the HPG axis and responds to offspring cues will lay the groundwork to explore any potential role this gene may play in reproductive physiology or behavior.

Rock doves (*Columba livia*) provide a powerful model to explore the dynamics of prolactin and its receptor across parental care and between the sexes. These birds form monogamous bonds and exhibit biparental care, with both sexes incubating eggs and provisioning offspring. Additionally, rock doves produce “crop milk” to feed their offspring, which is regulated by circulating prolactin (Abs, 1983; Horseman and Buntin, 1995). Unlike mammals, both male and female rock doves pseudo-lactate, allowing the comparison of sex differences without the confounds of pregnancy and female-only lactation. In doves, prolactin maintains incubation behaviors and facilitates the onset of chick provisioning, rising mid-incubation and peaking around hatch in both sexes (Austin et al., *in review*; Cheng and Burke, 1983; Horseman and Buntin, 1995; Ramsey et al., 1985). Prolactin then remains elevated post-hatching to facilitate both crop milk production and chick brooding/provisioning, a pattern typical of avian species with altricial young (Angelier and Chastel, 2009; Smiley, 2019). Additionally, we detected *PRLR* and *PRL* gene transcripts across the HPG axis in previous RNAseq studies (Austin et al., 2021a; Calisi et al., 2018; MacManes et al., 2017), setting a foundation to examine patterns of expression in these genes during parental care.

In this study, we examined how reproductive tissues vary in prolactin responsiveness and local prolactin expression across breeding and in response to offspring presence. Our goal was to understand how regulation at the tissue level may facilitate and coordinate reproductive transitions beyond circulating hormones alone. First, we characterized the expression of prolactin (*PRL*) and its receptor (*PRLR*) across multiple stages of parental care in the hypothalamus, pituitary, and gonads of both sexes. We also characterized these genes in the crop sac (“crop”), which is where crop milk is produced in doves. Then, we tested the influence of offspring cues on *PRL* and *PRLR* by introducing chicks at mid-incubation, before plasma prolactin is elevated and crops are fully functional for chick provisioning and lactation (Dong et al., 2012; Horseman and Buntin, 1995). We compared this “early hatch” manipulation to the equivalent stage at mid-incubation as a control group. Through this manipulation, we assessed to what degree offspring presence influences prolactin gene dynamics separate from the rise in circulating prolactin normally seen before hatch (Austin et al., 2021b). We hypothesized that offspring presence drives prolactin and prolactin responsiveness in key tissues. Therefore, we predicted plasma prolactin levels and *PRLR* expression would increase when chicks were added mid-incubation to compensate for normally low circulating prolactin levels at this stage. Alternatively, the priming effect of circulating prolactin before hatch may drive tissue responsiveness to prolactin. In this case, we predicted that chick presence alone would not increase *PRLR* expression, as hormonal priming was not yet completed. These hypotheses are not mutually exclusive and may be supported in some tissues under examination, but not others. Lastly, because both male and female rock doves exhibit the same suite of parental behaviors, we hypothesized that prolactin gene dynamics would be similar between the sexes.

## 2. Methods

This project was conducted in conjunction with a larger RNAseq study of the HPG axis during reproduction and parental care in rock doves (*Columba livia*). However, the focus of this study is prolactin-related gene dynamics in key tissues, including the crop. Thus, in addition to the HPG tissues (*n* ≅ 10/sex/sampling point, see <u>Supplemental Table 1</u> for exact sample sizes), we also collected crop tissue from a randomly-selected subset (n = 73) of these male-female pairs of breeding rock doves at focal stages of reproduction. We also collected crop tissues from an additional 20 individuals who were not part of the RNAseq study to increase sample sizes per stage (total n = 93, see <u>Supp. Table 1</u>). We focused on the following stages of reproduction: nest building (building), clutch completion/early incubation (incubation day 3: incubation begins when the first egg is laid in this species) (Abs, 1983), mid-incubation (incubation day 9), and the day the first chick hatched (hatch) (see Austin et al., 2021b, for more details). Additionally, to understand the influence of external cues on candidate gene expression, we also included a manipulation group (early hatch), where we experimentally reduced the length of the incubation period by replacing eggs with one young chick at mid-incubation (on incubation day 8) and then collected the pair ∼24 hours later. Circulating hormone data for these same individual birds across multiple stages of parental care were reported previously in Austin et al. (2021b). Here, we extend that study with the first gene expression data from these individuals, reporting *PRL* and *PRLR* gene counts across the hypothalamus, pituitary, and gonads and crop.

### 2.1 Study Animals

Rock doves *(Columba livia)* were socially housed in outdoor flight aviaries (1.5 x 1.2 x 2.1 m), each containing 8-10 breeding pairs, and were provided with nesting material (straw) and nest sites (wooden nest boxes, 16 per aviary). These outdoor aviaries exposed the birds to natural photoperiod for the area (Davis, California, USA), and photoperiod was supplemented with 14L:10D artificial lighting year-round. Birds were fed whole corn, turkey/game bird starter (30% protein; Modesto Milling, CA) and grit *ad libitum.* We used birds that were reproductively experienced and < 2 years old in this study. Further details can be found in Austin et al. (2021b).

### 2.2 Tissue Collection

Brain, pituitary, gonads, crop and trunk blood (for circulating hormones) were collected from birds at each timepoint following approved IACUC protocols (UC Davis protocol #20618). Tissues were flash frozen (brain, crop) or immediately placed in RNALater (Thermo Fisher) then flash frozen (pituitary and gonads) and stored at -80□ until use in downstream analyses. An additional 20 birds were collected in the same manner for crop tissues. For detailed collection methods and handling of HPG tissues (see Austin et al. 2021a; Calisi et al. 2018; MacManes et al. 2017), and for experimental design see Austin et al. (2021b). All of the subjects in this study, with the exception of the additional 20 birds collected for crop tissues alone, are included in Austin et al. (2021b).

### 2.3 RNA-sequencing for total gene expression

Before RNA processing, the hypothalamus and lateral septum were isolated using punch biopsy on a Leica CM 1860 cryostat and stored in RNALater at - 80 C before analysis (see Calisi et al 2018; MacManes et al, 2017 Austin et al. 2021a for details). Processing of brains, pituitaries, and gonads for RNA sequencing is described in detail in Austin et al., 2021a and Lang et al., 2020. Briefly, RNA from the hypothalamus, pituitary, and gonads was prepared for Illumina sequencing using the NEB Next Ultra Directional RNA Library Prep Kit, and sequenced on an Illumina HiSeq 400 via 125 base pair paired-end sequencing (Novogene). Reads were pseudomapped (*kallisto:* Bray et al., 2016) to the Rock Dove transcriptome v1.1.0 whose transcripts were annotated with genes from *Gallus gallus* genome v5 using BLAST. Transcriptomic data were then imported into the R statistical language using tximport (Soneson et al., 2016) and gene counts were variance-stabilized using the DEseq2 package (Love et al., 2014). Variance-stabilized gene counts for each sample were used in statistical analysis.

### 2.4 Quantitative PCR

To measure gene expression in the crop, we ran quantitative PCR (qPCR) on a subset of crops from each of the reproductive timepoints. For crop sample sizes by stage and sex, see <u>Supplemental Table 1</u>.

To extract total RNA from crops, we first homogenized an approximately 10 mg sample from each crop tissue using the OmniTip Tissue Homogenizer (Omni International), followed by RNA extraction using the Direct-zol RNA Miniprep kit (Zymo) with modifications recommended for lipid-rich tissues. We verified RNA purity and concentration using a NanoDrop 2000c (Thermo Scientific). For each sample, we treated 500 ng of RNA with DNase (Perfecta; QuantaBio) then performed cDNA reverse transcription using the QuantiTect Reverse Transcription Kit (Qiagen). We then ran real-time qPCR reactions with SYBR Green detection chemistry using the following reaction mix: 10 µL total reaction volume containing 1 µL cDNA template (diluted 1:5), 5 µL 2X SSOAdvanced SYBR Green PCR mix (BioRad), and 10 µM each of primer. We ran each reaction under the following conditions: 50□ for 2 min, 95□ for 10 min, and then 40 cycles of 95□ for 15 sec and 60□ for 30 sec. We ran samples in duplicate for each gene on the same 384-well plate using a CFX384 Touch Real-time PCR detection system (BioRad). We validated all primers for this study by running a 10-fold serial dilution to determine amplification efficiencies (average: 97.2% ± 5.63) and checked melt curves for a single product. Primer sequences, efficiencies, and amplicon lengths can be found in <u>Supplemental Table 2</u>.

We then quantified the relative expression of each gene of interest (*PRL* and *PRLR*) relative to the geometric mean of the reference genes, beta-actin (*ACTB*) and ribosomal protein L4 (*rpL4*) (Zinzow-Kramer et al., 2014) using the ddCt method (Livak and Schmittgen, 2001). We found no significant effect of reproductive stage (*F*_4,83_ = 1.6, *p* = 0.17), sex (*F*_1,83_ = 0.9, *p* =0.36) or their interaction (*F*_4,83_ = 1.2, *p* = 0.33) on mean reference gene expression, indicating stable reference genes for crop tissue. Samples that did not cross the cycle threshold within 40 cycles had Ct values set to 40. Normalized expression (dCt) was calculated as the average Cq value between technical replicates of each gene minus the geometric mean of the reference genes for each sample. We calculated relative expression (ddCt) as the normalized value (dCt) minus the average normalized expression for the nest-building stage. Nest-building was used as a reference as it was the first reproductive stage included in the study, and birds were not yet caring for eggs or chicks. Fold change equals 2^(-ddCt)^. We then log-transformed (log_e_ or ln) fold change values for statistical analysis to improve model fit and visualization.

### 2.5 Hormone measurements

Plasma hormones, including prolactin, were measured and described in rock doves across multiple stages of parental care in Austin et al., (2021b). Here, we used circulating prolactin data from Austin et al. (2021b) for our stages of interest (nest building, incubation day 3, incubation day 9, hatch and the manipulation on incubation day 8) to correlate plasma prolactin with *PRL* and *PRLR* gene expression, newly reported here. Briefly, plasma prolactin levels were measured from trunk blood using a heterologous radioimmunoassay (RIA) run at the Center for Biological Studies at Chizé, France (CEBC-CNRS) as detailed in (Angelier et al., 2007). This RIA had previously been validated in rock doves by creating a dose-response curve with pooled rock dove plasma and determining parallelism with standard curves consisting of chicken prolactin (Angelier et al., 2016a). Samples for this project were run in two separate assays with intra- and inter-assay coefficients of variation (CVs) of 9.58 and 11.83%, respectively. The minimal detectable prolactin level was 0.45 ng/ml.

### 2.6 Statistical analysis

All statistical analyses were performed in R (v.4.0.3, R Core Team, 2020). We compared gene expression in each gene-by-tissue combination using general linear models (glm), where gene expression (either variance-stabilized gene counts for RNAseq data or log-transformed fold change for qPCR) was predicted by stage, sex, and their interaction. We analyzed each gene-by-tissue combination in a separate model for three main reasons. First, we used two different methods for estimating gene expression, RNAseq and qPCR, and thus the expression data are not directly comparable across tissues. Second, we were interested in how each gene expression in tissue changed over time, responded to external manipulation, and varied by sex. Third, evidence shows that in different tissues genes for prolactin and its receptor are regulated by different promoters and transcription factors (Aoki et al., 2019; Featherstone et al., 2012), and therefore their expression should be considered independently.

For each glm, we ensured that our data met the model assumptions. If main effects were significant (alpha = 0.05), we compared group differences using pairwise comparisons of our *a-priori* hypotheses. The interaction between stage and sex was not significant for *PRL* and *PRLR* in any tissue, which suggests that males and females responded similarly across stage and to external manipulation. Because sex interactions were not significant, we did not include this term in future models (gene expression ∼ stage + sex). We also present estimates,standard errors, and *p*-values of *a priori* contrasts of biological interest. Following Austin et al. (2021b), we compared each reproductive stage to the adjacent or subsequent stage in the normal course of parental care: nest building vs. clutch completion, clutch completion vs. mid-incubation, and mid-incubation vs. hatch. This approach allowed us to compare gene expression changes during key reproductive transitions. We also compared how external manipulation affected gene expression, by comparing the early hatch group to its equivalent control stage, i.e., early hatch vs. mid-incubation. To determine if adding chicks mid-incubation had a similar effect to that seen when chicks naturally hatch after 18 days of incubation, we also compared early hatch vs. hatch. A list of pairwise contrasts can be found in Table 1. Finally, we examined relationships between plasma prolactin levels and gene expression within each tissue by calculating Spearman’s correlation coefficients (⍰).

**Table 1.**
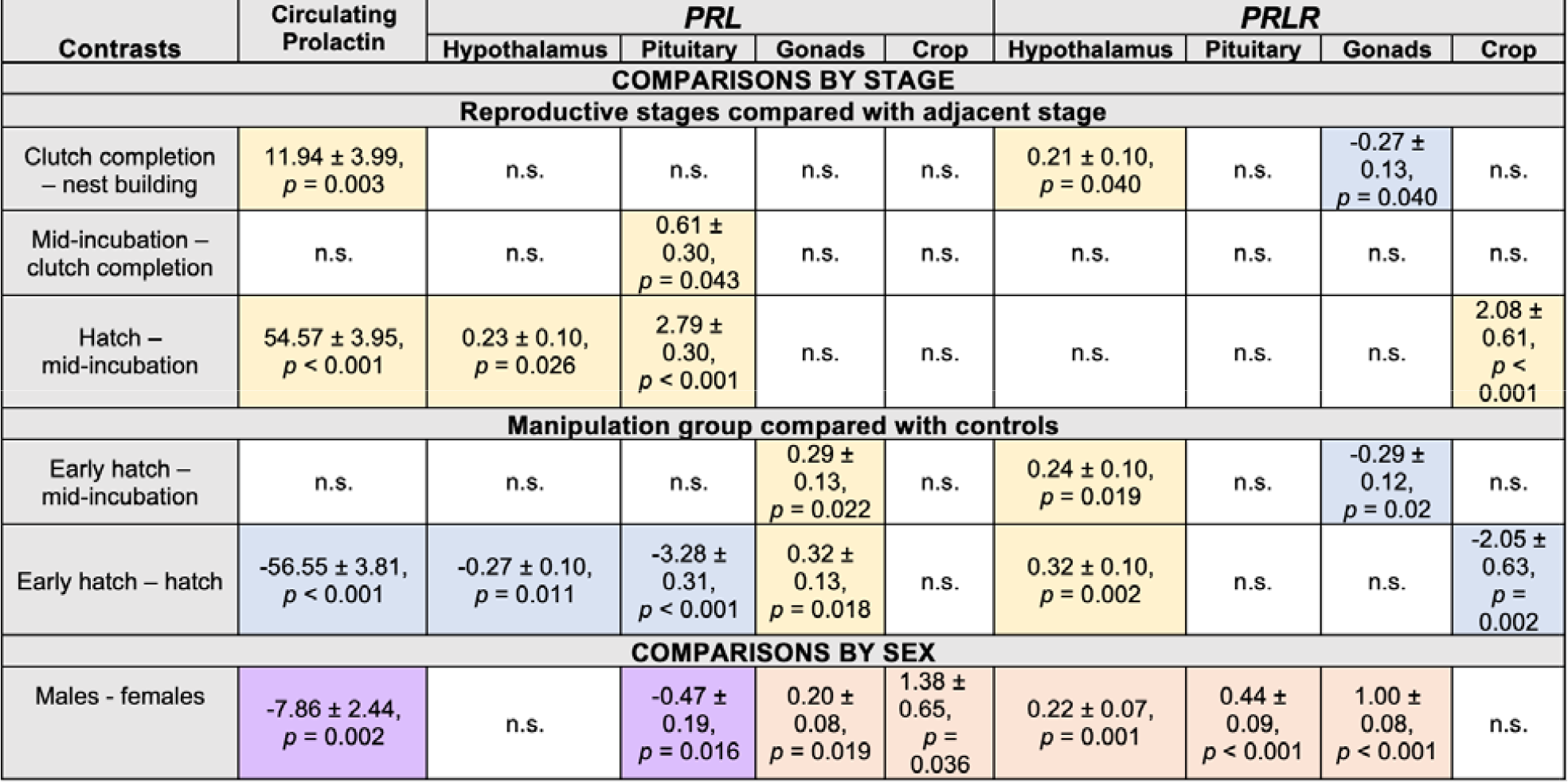
Pairwise contrasts for circulating prolactin, and *PRL* and *PRLR* within each tissue. Using *a priori* hypotheses, we developed contrasts to compare relevant transitions during parental care. We compared adjacent stages of reproduction and then compared the early hatch manipulation group with its equivalent control at mid-incubation and with concentration (circulating proalctin) /gene counts (*PRL* and *PRLR*) typically seen at natural hatch after 18 days of incubation. We also compared values between the sexes. Estimates ± standard errors are presented as A - B, where the estimate is group A minus group B. Only contrasts with *p*-values < 0.05 are shown. Comparisons where values increased in A relative to B are highlighted in yellow, where those where values decreased are in blue. For sex differences, purple indicates when values are higher in females and orange when values are higher in males.

| Contrasts | Circulating Prolactin | PRL |  |  |  | PRLR |  |  |  |
| --- | --- | --- | --- | --- | --- | --- | --- | --- | --- |
|  |  | Hypothalamus | Pituitary | Gonads | Crop | Hypothalamus | Pituitary | Gonads | Crop |
| COMPARISONS BY STAGE |  |  |  |  |  |  |  |  |  |
| Reproductive stages compared with adjacent stage |  |  |  |  |  |  |  |  |  |
| Clutch completion – nest building | 11.94 ± 3.99, *p* = 0.003 | n.s. | n.s. | n.s. | n.s. | 0.21 ± 0.10, *p* = 0.040 | n.s. | -0.27 ± 0.13, *p* = 0.040 | n.s. |
| Mid-incubation – clutch completion | n.s. | n.s. | 0.61 ± 0.30, *p* = 0.043 | n.s. | n.s. | n.s. | n.s. | n.s. | n.s. |
| Hatch – mid-incubation | 54.57 ± 3.95, *p* < 0.001 | 0.23 ± 0.10, *p* = 0.026 | 2.79 ± 0.30, *p* < 0.001 | n.s. | n.s. | n.s. | n.s. | n.s. | 2.08 ± 0.61, *p* < 0.001 |
| Manipulation group compared with controls |  |  |  |  |  |  |  |  |  |
| Early hatch – mid-incubation | n.s. | n.s. | n.s. | 0.29 ± 0.13, *p* = 0.022 | n.s. | 0.24 ± 0.10, *p* = 0.019 | n.s. | -0.29 ± 0.12, *p* = 0.02 | n.s. |
| Early hatch – hatch | -56.55 ± 3.81, *p* < 0.001 | -0.27 ± 0.10, *p* = 0.011 | -3.28 ± 0.31, *p* < 0.001 | 0.32 ± 0.13, *p* = 0.018 | n.s. | 0.32 ± 0.10, *p* = 0.002 | n.s. | n.s. | -2.05 ± 0.63, *p* = 0.002 |
| COMPARISONS BY SEX |  |  |  |  |  |  |  |  |  |
| Males - females | -7.86 ± 2.44, *p* = 0.002 | n.s. | -0.47 ± 0.19, *p* = 0.016 | 0.20 ± 0.08, *p* = 0.019 | 1.38 ± 0.65, *p* = 0.036 | 0.22 ± 0.07, *p* = 0.001 | 0.44 ± 0.09, *p* < 0.001 | 1.00 ± 0.08, *p* < 0.001 | n.s. |

## 3. Results

We examined the effect of reproductive stage and sex on plasma prolactin, and gene expression of *PRL* and *PRLR* in HPG and crop tissues. Results from *a priori* pairwise comparisons for all tissues and circulating prolactin can be found in <u>Table 1</u>.

### 3.1 Plasma prolactin levels

As in our larger analysis of circulating prolactin (Austin et al., 2021b), we found that plasma prolactin levels varied significantly across the stages examined in this study (stage: *F*_4,106_ = 83.6, p < 0.01) and with sex (*F*_1,106_ = 10.4, p < 0.01). Prolactin significantly increased from nest building to clutch completion, and from mid-incubation to hatch, but did not differ from clutch completion to mid-incubation (Fig.2). When chicks were added mid-incubation (early hatch), circulating prolactin did not significantly differ from the equivalent stage at mid-incubation, and was significantly lower than the level seen at typical hatching. Across all stages, females had significantly higher prolactin levels than males.

**Figure 1.**
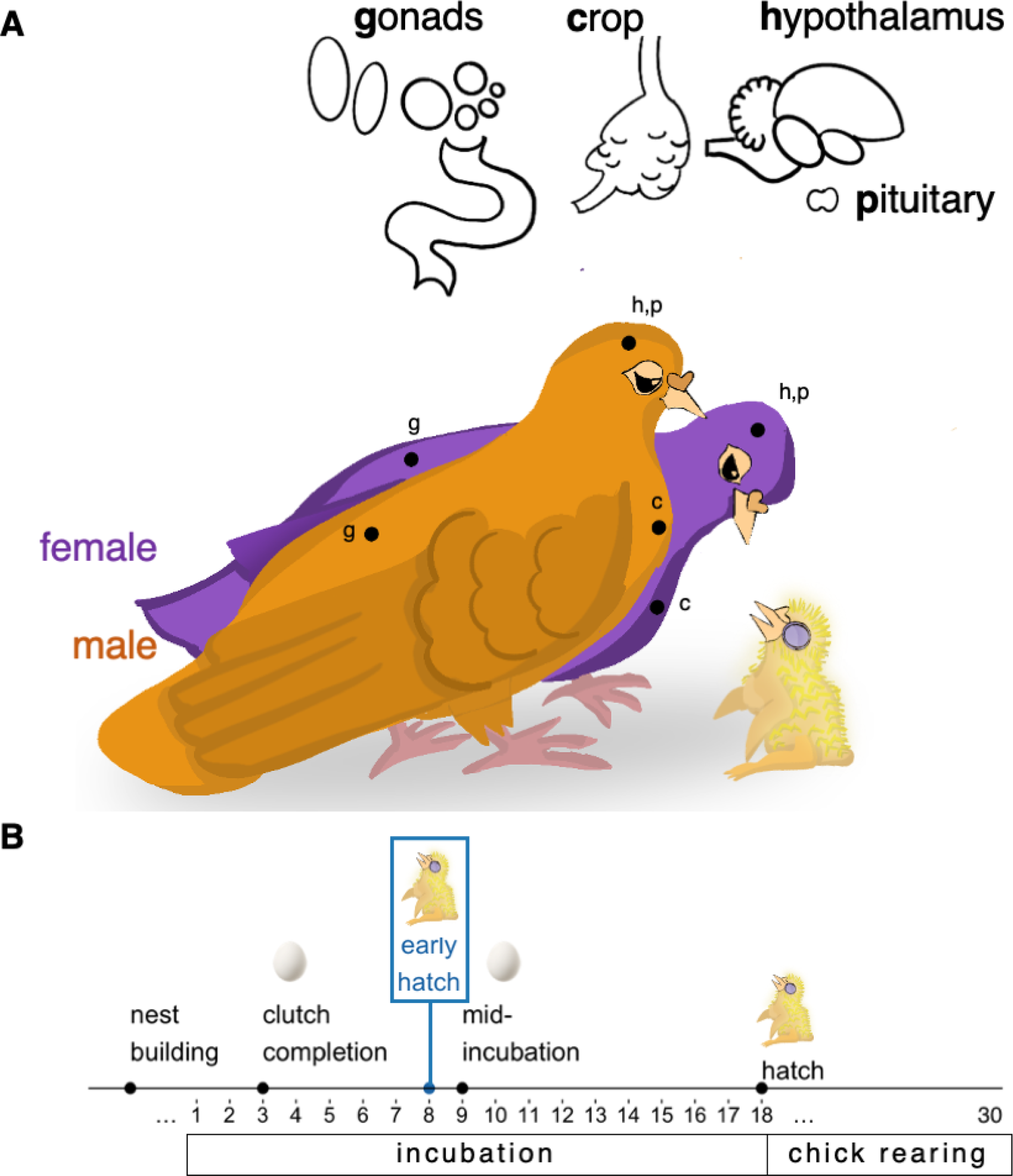
Schematic diagram of experimental design. (**A**) Tissues sampled in both males and females include the hypothalamus, pituitary, gonads (testes in males, ovaries and oviduct in females), and crop. Relative locations of each tissue are shown on the diagram with lowercase letters representing each tissue. (**B**) These tissues were collected from breeding pairs at the following stages of the rock dove reproduction: **nest building**, where pairs were engaged in nest building behaviors but had not yet laid an egg; **clutch completion** (incubation day 3), three days after the first egg was laid and the onset of incubation, when the second egg is laid (completing the two-egg clutch; this population of rock doves had a one day gap between laying the 1st and 2nd eggs); **mid-incubation** (incubation day 9), nine days after the first egg was laid and the onset of incubation; **hatch**, the day of the first chick hatching; and **early hatch**, a manipulation group where eggs were removed on the eighth day of incubation and replaced with a young chick(s) to test the impact of external cues (offspring presence) on gene expression and circulating prolactin concentration.

**Figure 2.**
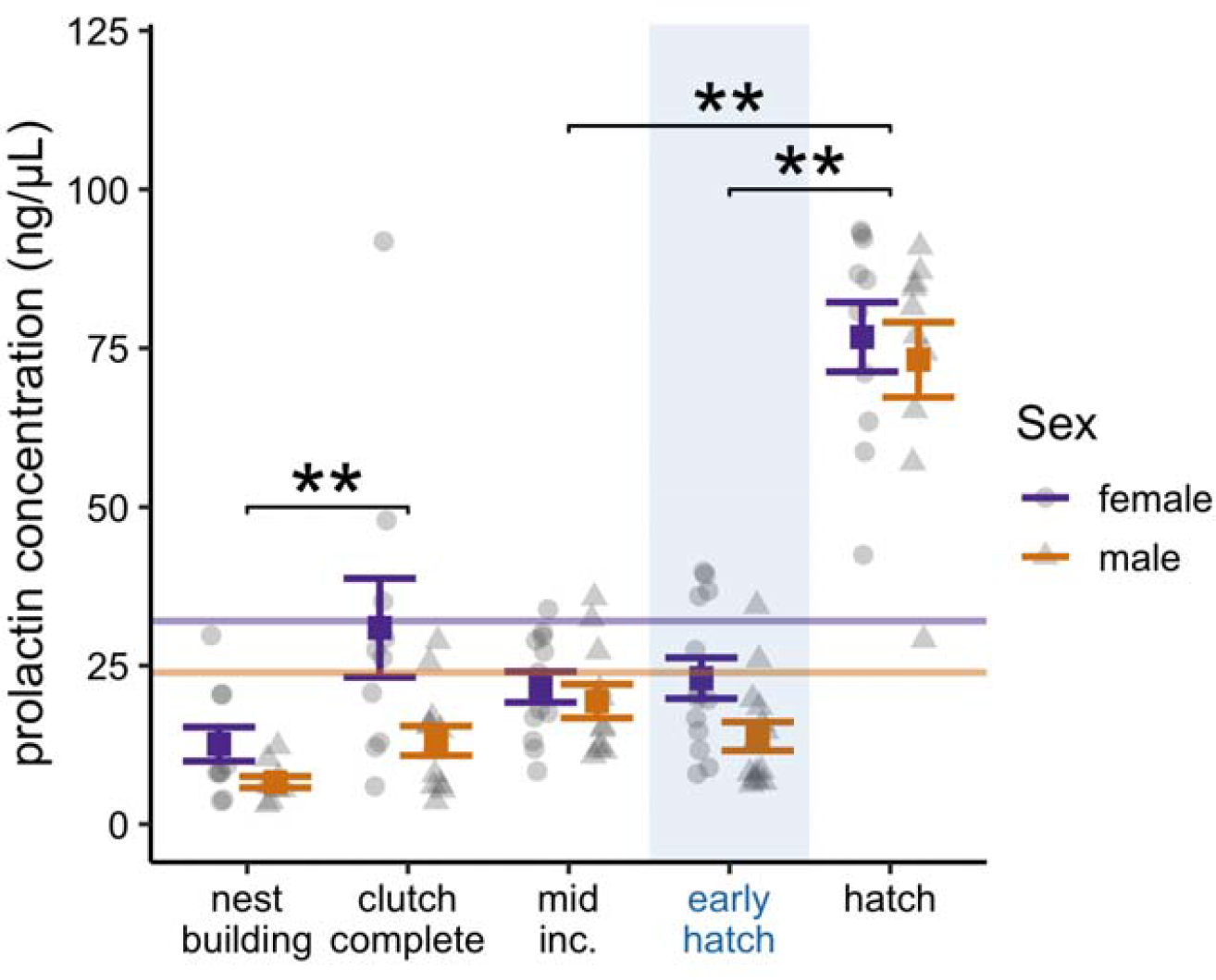
Plasma prolactin across reproductive stages. Prolactin plasma concentrations (ng/mL) of each stage for females (purple, triangles) and males (orange, circles). Means and standard errors are shown for each stage and sex. The mean value for each sex is shown as a horizontal line. Significant pairwise comparisons between stages are indicated (** *p* < 0.01; for a full list of *a priori* defined comparisons, see Table 1). Plasma prolactin data were originally presented in Austin et al., *in review*.

### 3.2 Hypothalamic PRL and PRLR expression

While there was no significant difference in hypothalamic *PRL* expression across stage in our models (*F*_4,94_ = 0.7, *p* = 0.569), this effect was largely driven by earlier time points. When we investigated a priori hypotheses of gene expression difference across stage, we found that birds at hatch had higher *PRL* expression compared with those at mid-incubation. When we investigated how external manipulation influenced gene expression, we found that the addition of chicks (early hatch) at mid-incubation did not significantly affect hypothalamic *PRL* levels above those seen at its control at mid-incubation. We found that *PRL* at early hatch was significantly lower than at a typical hatch (Fig. 3A, Table 1). We found a suggestive trend (0.05 < *p* < 0.10) of sex on hypothalamic *PRL* in our models (*F*_1,94_ = 2.9, *p* = 0.092), suggesting that males may express hypothalamic *PRL* at slightly higher levels than females. Hypothalamic *PRL* and plasma prolactin levels were not significantly correlated (Fig.4A; ⍰*_99_* = 0.12, *p* = 0.200).

**Figure 3.**
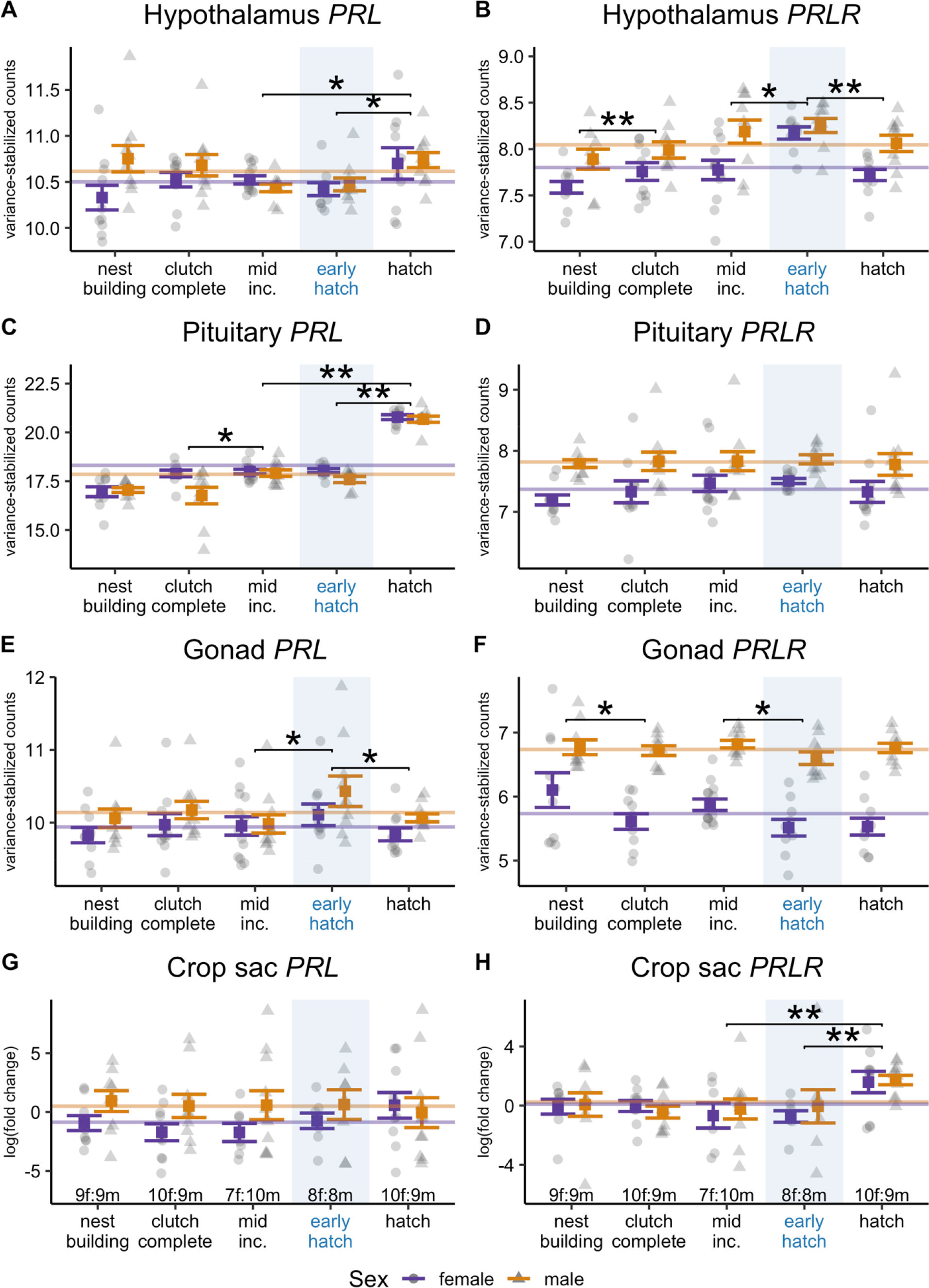
*PRL and PRLR* gene expression across tissues. *PRL* (left) and *PRLR (*right) expression, respectively, in the (**A-B**) hypothalamus, (**C-D**) pituitary, (**E-F**) gonads (testes and ovaries /oviducts), and (**G-H**) crop across reproductive stages. Early hatch, a manipulation group where we added chick(s) at mid-incubation, is highlighted in blue. Male (orange, triangles) and female (purple, circles) means and standard errors of the gene count mean (SEM) for each stage. The mean value for each sex is shown as a horizontal line. Significant pairwise comparisons between stages are indicated (* *p* < 0.05, ** *p* < 0.01; for a full list of *a priori* contrasts, see Table 1).

**Figure 4.**
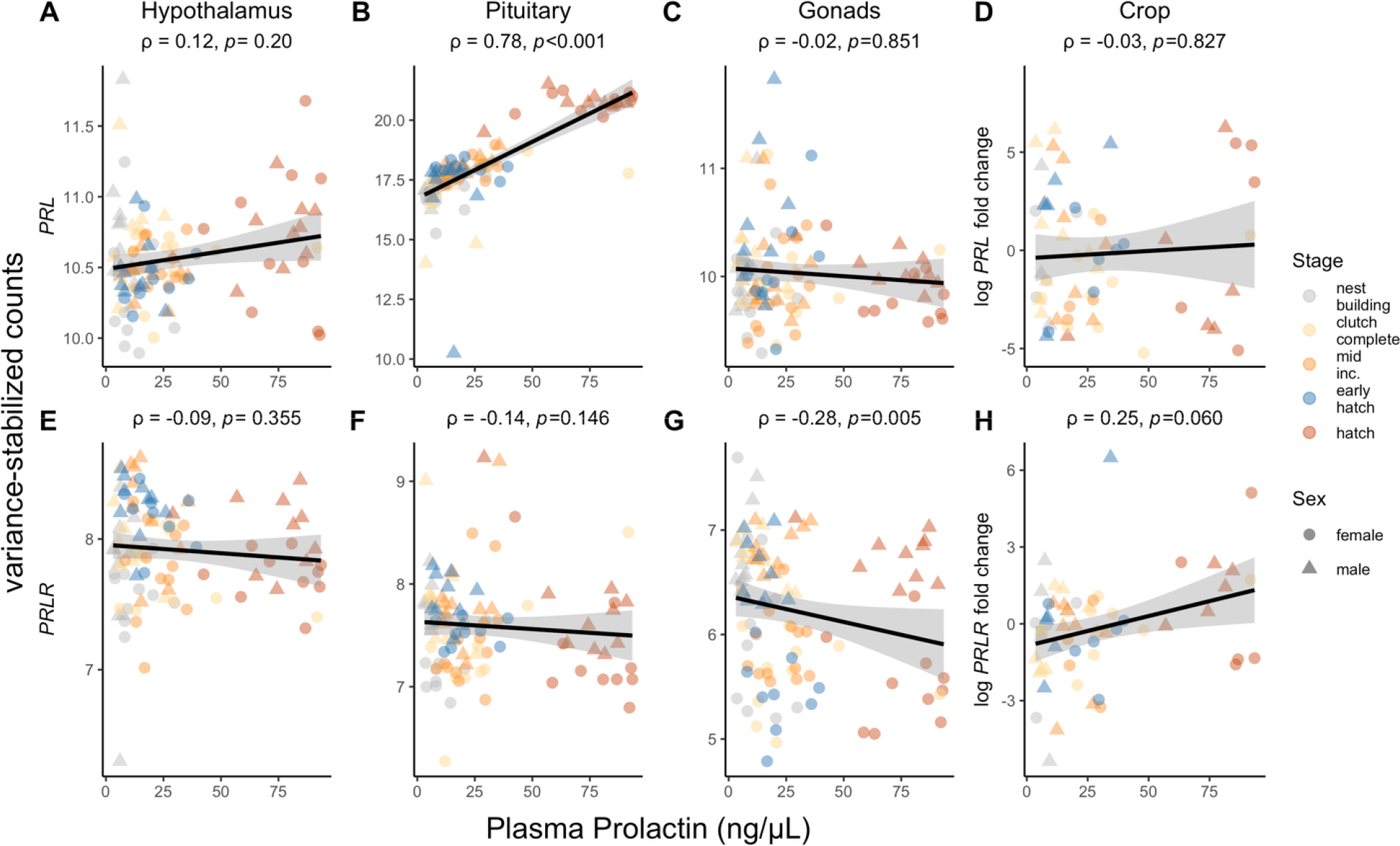
Correlations between plasma prolactin concentrations and gene expression across tissues. Correlations between plasma prolactin (as measured by RIA) and gene expression of (**A**) hypothalamic *PRL*, (**B**) pituitary *PRL*, (**C**) gonadal *PRL*, (**D**) crop *PRL*, (**E**) hypothalamic *PRLR*, (**F**) pituitary *PRLR*, (**G**) gonad *PRL*, and (**H**) crop *PRLR* for each individual bird. Spearman’s correlation coefficient (⍰) and *p*-values are shown for each gene-tissue combination. Gray shading around the line of best fit represents the 95% confidence interval. Point color corresponds to reproductive stage, and males and females are indicated with circles and triangles, respectively.

Hypothalamic *PRLR* expression significantly differed by stage (*F*_4,94_= 7.7, *p* < 0.01) and sex (*F*_1,94_ = 10.8, *p* < 0.01). Specifically, *PRLR* counts increased at clutch completion compared with nest building (Fig. 3B; Table 1). When we compared the early hatch manipulation to its equivalent control at mid-incubation, *PRLR* levels significantly increased (Fig. 3B, Table 1). Further, *PRLR* expression at the early hatch manipulation was also significantly higher compared to hatch. We found no significant correlation between hypothalamic *PRLR* and plasma prolactin (Fig.4E; ⍰*_99_* = -0.09, *p* = 0.355).

### 3.3 Pituitary PRL and PRLR expression

Like plasma prolactin, pituitary *PRL* gene expression varied significantly with stage (*F*_4,98_ = 47.9, *p* < 0.001) and sex (*F*_1,98_ = 6.0, *p =* 0.016). Pituitary *PRL* expression also increased from mid-incubation to hatching (Fig. 3C; Table 1). Unlike plasma prolactin levels, however, pituitary *PRL* significantly increased from clutch completion to mid-incubation but did not significantly change from nest building to clutch completion (Table 1). Pituitary *PRL* gene counts were significantly higher in females than males, as seen in plasma levels. As expected, pituitary PRL expression and plasma prolactin were significantly positively correlated (Fig.4B; ⍰*_101_* = 0.78, *p* < 0.001).

Pituitary *PRLR*, in contrast, did not significantly differ across stages (Fig. 3D; *F*_4,98_ = 0.7, *p* = 0.616). However, we found a significant effect of sex (*F*_1,98_ = 29.0, *p* < 0.001), where males expressed higher levels of pituitary *PRLR* than females. Unlike pituitary *PRL*, *PRLR* expression did not correlate with plasma prolactin levels (Fig 4F; ⍰*_101_* = -0.14, *p* = 0.146).

### 3.4 Gonadal PRL and PRLR expression

*PRL* expression in the testes and ovaries/oviducts did not significantly differ with stage, though there was a suggestive trend (*F*_4,98_ = 2.2, *p* = 0.079). This trend appears to be driven by the early hatch manipulation, which significantly increased gonadal *PRL* compared to the mid-incubation control and hatching stages (Fig. 3E, Table 1). Gonadal *PRL* also differed significantly by sex (*F*_1,98_ = 5.7, *p* = 0.019), where testes expressed *PRL* at higher levels than ovaries and oviducts. We found no correlation between gonadal *PRL* expression and plasma prolactin (Fig. 4C; ⍰*_101_* = -0.02, *p* = 0.851).

Gonadal *PRLR* expression significantly differed with stage (*F*_4,98_ = 3.0, *p* = 0.023). Gonadal *PRLR* decreased at clutch completion compared with nest building, but did not differ from clutch completion to mid-incubation or mid-incubation to hatch (Fig. 3F, Table 1). At early hatch, gonadal *PRLR* expression did not significantly change compared to mid-incubation levels, though early hatch levels were significantly lower than at hatch. Further, there was a significant sex effect (*F*_1,98_ = 154.4, p < 0.001), where testes expressed more *PRLR* than ovaries/oviducts at all stages. Gonadal *PRLR* expression was significantly negatively correlated with plasma prolactin (Fig.4G; ⍰*_101_* = -0.28, *p* = 0.005).

### 3.5 Crop PRL and PRLR expression

In the crop, *PRL* expression remained relatively constant, with no significant stage effect detected (Fig. 3G; *F*_4,83_ = 0.21, *p* = 0.930). However, we found crop *PRL* expression differed by sex (*F*_1,83_ = 4.50, *p* = 0.037) with males having higher *PRL* than females. Crop *PRL* was not correlated with plasma prolactin (Fig.4D; ⍰*_80_* = -0.03, *p* = 0.827).

Unlike *PRL*, crop *PRLR* expression differed significantly by stage (*F*_4,87_ = 4.30, *p* = 0.003). This effect was likely driven by increased expression at hatch, which was higher than every other stage in contrasts (Fig. 3H, Table 1). However, crop *PRLR* levels did not significantly differ after the early hatch manipulation compared to mid-incubation controls. We did not find a significant effect of sex on crop *PRLR* expression (*F*_1,87_ = 0.19, *p* = 0.665). We found a suggestive positive correlation between crop *PRLR* and plasma prolactin (⍰*_80_* = 0.25, *p* = 0.060), which is likely driven by levels at hatch (Fig.4H).

## 4. Discussion

We characterized how circulating prolactin, *PRL* and *PRLR* gene expression in the HPG axis and crop varied across four reproductive stages (nest building, clutch completion, mid-incubation, and hatch) in both male and female rock doves. We then tested how externally manipulating the development period by adding offspring halfway through incubation influenced prolactin and HPG and crop tissue *PRL* and *PRLR* gene expression levels ∼24 hours later. This study thus provides a finer resolution into how prolactin gene expression changes across specific reproductive stages and within specific tissues important for parental care.

We found that circulating prolactin was lowest at nest building and highest after chicks hatch. Pituitary *PRL* gene expression mirrored this pattern, as expected. Hypothalamic *PRL* also increased at hatching. We did not observe significant differences in gonad or crop *PRL* expression across the reproductive stages measured. *PRLR* expression also did not differ across reproductive stages in the HPG or crop. However, some tissues showed significant increases in *PRLR* across specific transitions during parental care (such as from nest building to early incubation), though the overall effect size of these increases was relatively small, and the biological significance of these changes remains to be tested. We also found significant sex differences in prolactin and *PRL/PRLR* gene expression. In response to offspring presence, we found no significant difference in circulating prolactin levels as compared to the mid-incubation control. However, chick presence significantly increased hypothalamic *PRLR* and decreased gonadal *PRL*. The early hatch manipulation did not affect pituitary or crop gene expression.

### 4.1 Characterization of PRL and PRLR expression across the HPG and crop

In the hypothalamus, a key regulatory center for reproductive and parental behavior, we found that *PRL* gene expression increased when chicks hatched. Prolactin can act upon hypothalamic nuclei, such as the preoptic area (POA), to regulate key parental behaviors in birds and mammals (Brown et al., 2017; Dobolyi et al., 2014; Slawski and Buntin, 1995). Prolactin can also physiologically coordinate parental care through actions on the hypothalamus, such as affecting overall HPG axis regulation via hypothalamic gonadotropin releasing hormone (GnRH) neurons (Grattan et al., 2007; Rozenboim et al., 1993), or regulating energy balance and hyperphagia through hypothalamic neuropeptide Y (Buntin et al., 1991; Lopez□ Vicchi et al., 2020; Slawski and Buntin, 1995). In birds, both prolactin protein and gene expression, as well as prolactin binding and receptors have been identified in the hypothalamus (Buntin and Ruzycki, 1987; Buntin and Walsh, 1988; Chaiseha et al., 2012; Ramesh et al., 2000; Smiley et al., 2021). We found that *PRL* expression significantly changed from mid-incubation to hatching in the brain. This is consistent with rodent studies, where hypothalamic *PRL* mRNA also increased from pregnancy to lactation in female rats (Torner et al., 2004, 2002). Extra-pituitary *PRL* may play a role in regulating the stress hyporesponsiveness seen during maternal care (Torner et al., 2004), though it remains unclear whether hypothalamic *PRL* is actually translated into a functional protein. We thus extend previous studies characterizing hypothalamic *PRL* expression in the avian brain by showing that its expression changes during parental care.

We also found that hypothalamic *PRLR* increased from nest building to clutch completion. This increase in hypothalamic responsiveness to prolactin may facilitate incubation behavior (Buntin 1996). Studies in birds have linked incubation behavior with increases in circulating prolactin (Angelier and Chastel, 2009; Hope et al., 2020; Ramsey et al., 1985; Sockman et al., 2000). We found that plasma prolactin increased from nest building to clutch completion (the third day of incubation in this species), and that hypothalamic *PRLR* also increased during this transition. This increase suggests that behavioral centers in the brain become more responsive to prolactin levels to facilitate incubation behavior. Previous studies show intracerebroventricular injections of prolactin increased incubation in turkey hens (Youngren et al 1991), but did not induce incubation in ring doves (Buntin and Tesch, 1985). In light of our findings, it is possible that the isolated, non-breeding doves in Buntin and Tesch (1985) may have not upregulated *PRLR* levels sufficiently to behaviorally respond to the injections of prolactin. Progesterone and estradiol may also upregulate hypothalamic *PRLR* during this transition, as these hormones facilitate incubation in doves (Michel, 1977; Silver, 1978). However, prolactin itself does not appear to upregulate its receptor in the hypothalamus, as we found no significant correlation between hypothalamic *PRLR* and plasma prolactin. This result differs from turkey hypothalamic *PRLR*, which was negatively correlated with plasma prolactin (Zhou et al., 1996). Causal studies which manipulate prolactin or other hormones involved in incubation behavior are needed to further understand drivers of hypothalamic *PRLR* across this transition.

In the pituitary, we found that *PRL* gene expression mirrored plasma prolactin patterns, while *PRLR* did not. Nearly all circulating prolactin originates from lactotroph cells in the anterior pituitary (Freeman et al., 2000), thus, a correlation between pituitary *PRL* and plasma prolactin was expected. Like plasma levels, pituitary *PRL* was lowest at nest building, rose at clutch completion/early incubation, and peaked at hatch, consistent with other studies in doves and pigeons (Cheng and Burke, 1983; Dong et al., 2012; Horseman and Buntin, 1995). Slight differences in pituitary *PRL* in comparison to plasma levels may be due to different drivers for prolactin peptide secretion versus gene transcription. For instance, we observed a significant increase in plasma prolactin at clutch completion, but no concordant significant change in pituitary *PRL.* Stored prolactin peptide may have been released to facilitate the onset of incubation, as prolactin has been shown to increase after the first egg is laid (when incubation begins in doves;(Cheng and Burke, 1983; Lea et al., 1986). Vasoactive intestinal peptide (VIP), a neuropeptide that stimulates the release of prolactin from the pituitary in birds (Macnamee et al., 1986; Vleck and Patrick, 1999) typically increases around incubation as well (Cloues et al., 1990) which may have caused prolactin release but not upregulation of *PRL*. Later, we find that pituitary *PRL* mRNA significantly increased from clutch completion to mid-incubation, but observed no change in plasma levels. This difference may be due to increased lactotroph recruitment in the pituitary (Pitts et al., 1994), which would lead to higher overall *PRL* transcription that may be stored for release later in incubation. Lastly, we found that *PRLR* did not change across the reproductive stages measured. While the role of the *PRLR* in the pituitary remains unclear, it may play a role in autocrine negative feedback (as seen in mammals; Ferraris et al., 2012). However, this potential role remains untested in birds.

Gonadal *PRL* and *PRLR*, in contrast, did not differ across the reproductive stages we measured. We found no significant changes in gonadal *PRL* in either sex, though gonad *PRLR* increased in both the testes and ovaries/oviducts at clutch completion compared to nest building. Prolactin treatment has been shown to have an anti-gonadal effect in birds, leading to reduced gonad size (Meier, 1969; Meier et al., 1971) and sex steroid secretion (Camper and Burke, 1977; Reddy et al., 2002). In chickens, FSH, but not LH, increased ovarian *PRLR* (Hu et al., 2017). However, FSH has been shown to increase during nest building and decrease around ovulation / laying in doves (Cheng and Balthazart, 1982), which does not support that FSH may drive gonadal *PRLR*. This relationship, however, may differ across sexes and species. In male rats, for instance, FSH treatment decreased testicular *PRLR* expression in the Sertoli cells (Guillaumot and Benahmed, 1999). *PRLR* may play a role in spermatogenesis, as hyperprolactinemia reduces sperm count in mammals (Gill-Sharma, 2009), though such a relationship remains unstudied in birds. The increased gonadal *PRLR* in this study could indicate that *PRLR* regulates sex steroids, which are often higher before laying than during parental care in doves (Austin et al., 2021b; Dong et al., 2012; Feder et al., 1977). We did not find significant changes in estradiol or testosterone between nest building and clutch completion (where gonadal *PRLR* increased) (Austin et al., 2021b), though progesterone levels fluctuate significantly as birds began incubation (Austin et al., 2021b). Increased prolactin responsiveness in the gonads may possibly alter steroidogenic pathways and progesterone release, though this has not been tested. Clearly, more comparative research into the effects of prolactin on gametogenesis and steroidogenesis in the gonads is needed.

In the crop, *PRLR* expression patterns more closely mirrored plasma prolactin levels, though we found no variation in crop *PRL*. Like circulating prolactin and pituitary *PRL*, crop *PRLR* significantly increased at hatching, but did not differ across nest building and incubation. This pattern is consistent with crop weight changes across the dove breeding cycle, where crop thickness and weight peaks around hatching in conjunction with crop milk production (Cheng and Burke, 1983). As the crop is highly responsive to prolactin (Horseman and Buntin, 1995; Riddle and Braucher, 1931) and prolactin regulates its own binding in this tissue (Shani et al., 1981), our results reiterate that prolactin levels likely drive crop *PRLR* gene expression. Crop *PRLR* dynamics are consistent with mammalian mammary gland cells, where prolactin also upregulates *PRLR* expression (Bera et al., 1994; Swaminathan et al., 2008). While low, relative expression of *PRL* was detectable in both sexes. In mammals, autocrine ePRL plays a role in mammary gland differentiation and initiation of lactation (Chen et al., 2012), as well as in milk protein expression (Hennighausen et al., 1997). Unlike the mammary gland, the crop epithelium proliferates but does not differentiate (Gillespie et al., 2013); whether autocrine *PRL* plays a role in crop development remains unknown. Our results show that prolactin gene dynamics may be similar across convergently evolved organs for lactation, which opens the door for exciting comparative studies of “milk” production across species.

### 4.2 Effects of offspring presence on PRL and PRLR gene expression

In response to the early hatch manipulation, where chicks were added mid-incubation to examine response to offspring presence, neither circulating/plasma prolactin levels nor pituitary *PRL* expression significantly changed compared to mid-incubation. Exposure to chicks increased plasma prolactin in previous studies (Buntin, 1979; Hansen, 1966; Lea and Vowles, 1985). In doves, chick exposure for four days in early or mid-incubation led to significant increases in crop weight, suggesting increased prolactin (as the crop is known to be highly prolactin-responsive) (Hansen, 1966). In parental doves deprived of their own young for 24 hours, pituitary reserves of prolactin decreased after just one hour of chick exposure, indicating prolactin was released into circulation from the pituitary (Buntin, 1979). However, we did not see an increase in plasma prolactin or pituitary PRL transcription after 24 hours of chick exposure. This lack of response may have occurred because our sampling time course (≅24 hours after hicks were added) may have missed the window of any significant changes in prolactin. We may have missed an initial spike in prolactin release or transcription, as Buntin (1979) observed after one hour with chicks. Alternatively, 24 hours may have been not enough time to reliably upregulate *PRL* transcription or release. Secondly, it is possible that sufficient priming, either by hormonal secretion or internal rhythms during incubation, had not occurred. Indeed, 5 hours of offspring presence only stimulated prolactin release in non-breeding female ring doves that had been primed through estradiol and progesterone treatments (Lea and Vowles, 1985). In previous studies, doves were already in a chick-rearing state (deprived of their own chicks; Buntin, 1979) or had been given sufficient time to respond (i.e., more than one day; Hansen, 1966). Thus, if the manipulation had occurred later in incubation and closer to a natural hatch date, birds may have been more flexible in their ability to elevate prolactin in response to chick cues. Comparisons of our findings with a manipulation later in incubation could test this hypothesis.

Although plasma prolactin remained unchanged, hypothalamic *PRLR* increased when chicks were added, to levels significantly above those of mid-incubation or typical hatch. This increase suggests that neural responsiveness to prolactin may have increased to compensate for the typically low circulating prolactin at this stage and to facilitate a parental response to chicks. Indeed, parental behaviors can spontaneously occur without subsequent increases in prolactin (Wang & Buntin 1999), and we observed parents brooding and attempting to feed chicks during this manipulation (Austin et al., 2021b). This behavior may have been facilitated by the increasing responsiveness to prolactin in hypothalamic nuclei like the POA, where prolactin is critical for chick feeding in doves (Buntin et al., 1991; Slawski and Buntin, 1995). Our findings also suggest that the hypothalamus may be able to respond more quickly to offspring cues than prolactin release from the pituitary, as plasma prolactin remained unchanged after the same period of chick exposure. Although we examined the hypothalamus as a whole, future examination of specific nuclei or cell-types could clarify where this *PRLR* response occurs.

We also observed significant upregulation of *PRL* and downregulation *PRLR* in the gonads of both sexes. Like hypothalamic *PRLR*, gonadal expression of these genes differed from both the mid-incubation control and typical hatch. Studies show that sex steroids like estradiol or progesterone are required to exhibit parental behaviors in birds (El Halawani et al., 1986; Hutchison, 1975), including response to chicks (Lea and Vowles, 1985). As previously suggested, increased local *PRL* transcription could shift steroidogenic pathways to increase necessary sex steroid production and facilitate a parental response. However, this hypothesis is not supported by the fact that estradiol significantly decreased in females in this study when chicks were added mid-incubation, and testosterone remained unaffected (Austin et al., 2021b). Alternatively, altered prolactin regulation could play a role in a gonadal stress response, as this manipulation increased circulating corticosterone in this study compared to mid-incubation (Austin et al., 2021b). This hypothesis is not supported because gonadal *PRL* or *PRLR* transcription did not change in non-breeding rock doves after an acute stressor (Calisi et al., 2018), though this response may differ when animals are in a parental state. Lastly, it is unclear why *PRLR* would be downregulated, ostensibly reducing prolactin responsiveness in the gonads. The gonadal response in *PRL* and *PRLR* could diverge because the two genes respond to different factors beyond prolactin, such as changes in other hormones or transcription factors that were affected during manipulation. These two genes do exhibit differential regulation in mammalian cells (Aoki et al., 2019; Featherstone et al., 2012), which if true in birds, could partially explain their opposing responses to chick presence. Overall, the gonadal transcriptional response to offspring cues merits further study to understand its importance in parental behavior.

### 4.3 Sex differences in PRL and PRLR gene expression

In almost all tissues examined, we uncovered consistent patterns of sex differences in *PRL* and *PRLR* gene expression. We found that females had higher levels of plasma prolactin and pituitary *PRL* expression than males, but males expressed higher levels of *PRLR* than females in all tissues. These sex differences are consistent with other studies in biparental birds, where females also had higher plasma prolactin than males (Hector & Goldsmith, 1985, Vleck 1998). In mammals, higher prolactin levels in females are explained by estrogen-responsive elements in the *PRL* gene promoter (Maurer and Notides, 1987), though this mechanism remains unconfirmed in birds (Kurima et al., 1995). Interestingly, we found no significant sex differences in hypothalamic *PRL*, and gonadal *PRL* was more expressed in males than females. This result is consistent with the idea that gene regulation differs for extra-pituitary *PRL* compared to “dogmatic” pituitary *PRL*, which has been found in mammalian cell lines (Marano and Ben-Jonathan, 2014). Further studies are needed to determine whether autocrine extra-pituitary prolactin could compensate locally for sex differences in circulating prolactin of pituitary origin. Our finding that males had higher *PRLR* across all tissues differs from rodent studies, where *PRLR* expression or prolactin-binding is often lower in males than females in the brain (Cabrera-Reyes et al., 2015; Pi and Voogt, 2002; Salais-López et al., 2018). The mechanism by which male doves may upregulate *PRLR* remains unclear, though testosterone may play a possible role, as castration significantly reduces prolactin binding in the rat brain (Salais-López et al., 2018). Our findings highlight the need to study prolactin dynamics in both sexes, as most studies of *PRL/R* expression in birds to date only included one sex or did not compare sex differences (Buntin and Buntin, 2014; Chaiseha et al., 2012; Ramesh et al., 2000; Smiley et al., 2021, 2020).

Together, our results support the hypothesis that different gene expression pathways can allow sexes to converge on a behavioral phenotype, preventing behavioral differences rather than promoting them (De Vries, 2004). A compensatory mechanism appears to be at play in our study, where females produced more hormonal signal (prolactin), but males increased downstream tissue responsiveness to that signal (via *PRLR*). This compensation may allow the sexes to exhibit a similar suite of parental behaviors despite sex differences in circulating prolactin levels. Indeed, several other bird species also exhibit higher prolactin levels in females, but both sexes show similar parental behaviors (Angelier et al., 2007; Angelier and Chastel, 2009). Sex differences in brain and peripheral *PRLR* may explain how similar parental behaviors can be maintained despite sex-biased differences in circulating prolactin. While much focus is on sex differences in behaviors or hormone levels, our results highlight the need to examine underlying mechanisms that may allow the sexes to converge to a similar phenotype (McCarthy and Konkle, 2005). Examining hormone and receptor dynamics in both sexes will be important to determine if this pattern occurs in other biparental species.

## 5. Conclusions

In summary, we report dynamic expression of prolactin and its receptor in various tissues important for reproduction and parental care, including the HPG endocrine axis and the crop. By examining specific stages of reproduction and parental care, we show that subtle changes in tissue-specific gene expression may help coordinate the overall response to prolactin and transitions between parental phenotypes. We show that *PRL* and *PRLR* gene expression in key tissues like the hypothalamus and gonads can respond to offspring cues even when plasma prolactin levels remain unaffected. Our results emphasize the need to examine how target tissues and endocrine axes transcriptomically respond to changing offspring stimuli, even in the absence of hormonal changes. Lastly, we uncovered consistent sex differences in prolactin regulation across the HPG axis, suggesting a compensatory mechanism by which the sexes may converge on similar parental behaviors in a biparental system. Future studies will be required to determine how regulation of these genes differs across tissues and the sexes, including manipulations of hormones that may drive gene expression. Overall, this study shows that tissue- and sex-specific changes in local production or responsiveness to a hormone can occur across an endocrine axis to coordinate physiological and behavioral breeding transitions.

## Acknowledgements

We thank T.P. Hahn and D. Caillaud for their helpful comments on earlier versions of this project. C. Parenteau also provided invaluable help with prolactin radioimmunoassays. We also thank the numerous undergraduate members of the Calisi laboratory for their dedication to animal care and husbandry: M. Alvarez, T. Chen, H. Hudson, E. Krestoff, C. Nguyen, A. Martinez, I. Orellana Bonilla, J. Sahota, E. Saldana, and D. Sanpedro. This project was funded by NSF IOS 1455960 to RMC and MDM, and a University of California Davis Dean’s Mentorship Award (2018) to BMNU and VSF. Comments from two anonymous reviewers greatly improved this manuscript.

---

**Supplemental Table 1.**
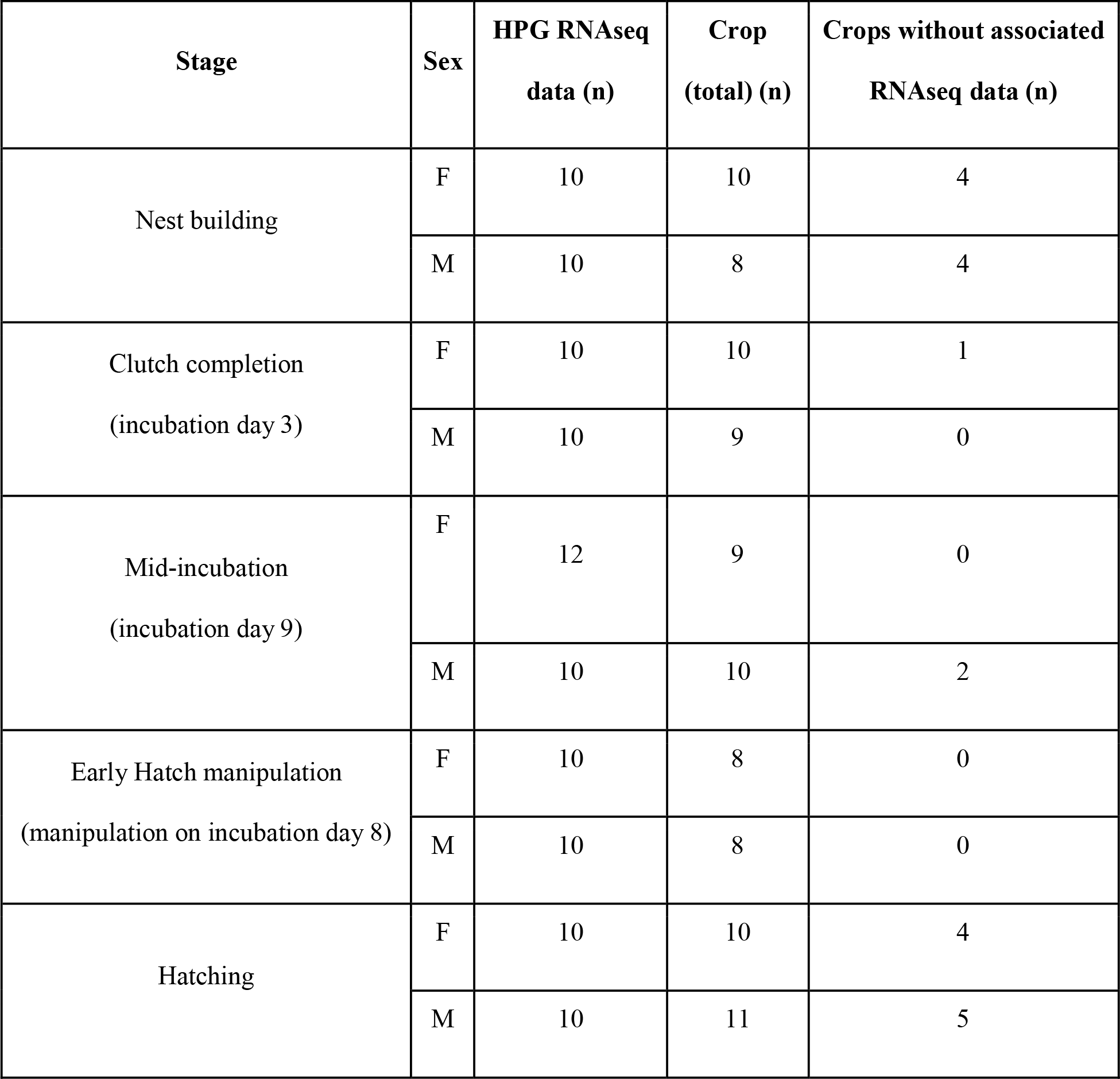
Sample sizes for tissues by stage and sex. Total sample sizes (n) are shown for the HPG RNAseq data (all individuals included had gene count data for all three tissues, the hypothalamus, pituitary and gonads). Total crop sample sizes are shown by stage and sex. The majority of crops came from individuals who also had HPG RNAseq data, except for 20 additional individuals that were included to increase crop sample size. The number of additional crops that were collected separately from the RNAseq study are shown in the right column.

**Supplemental Table 2.**
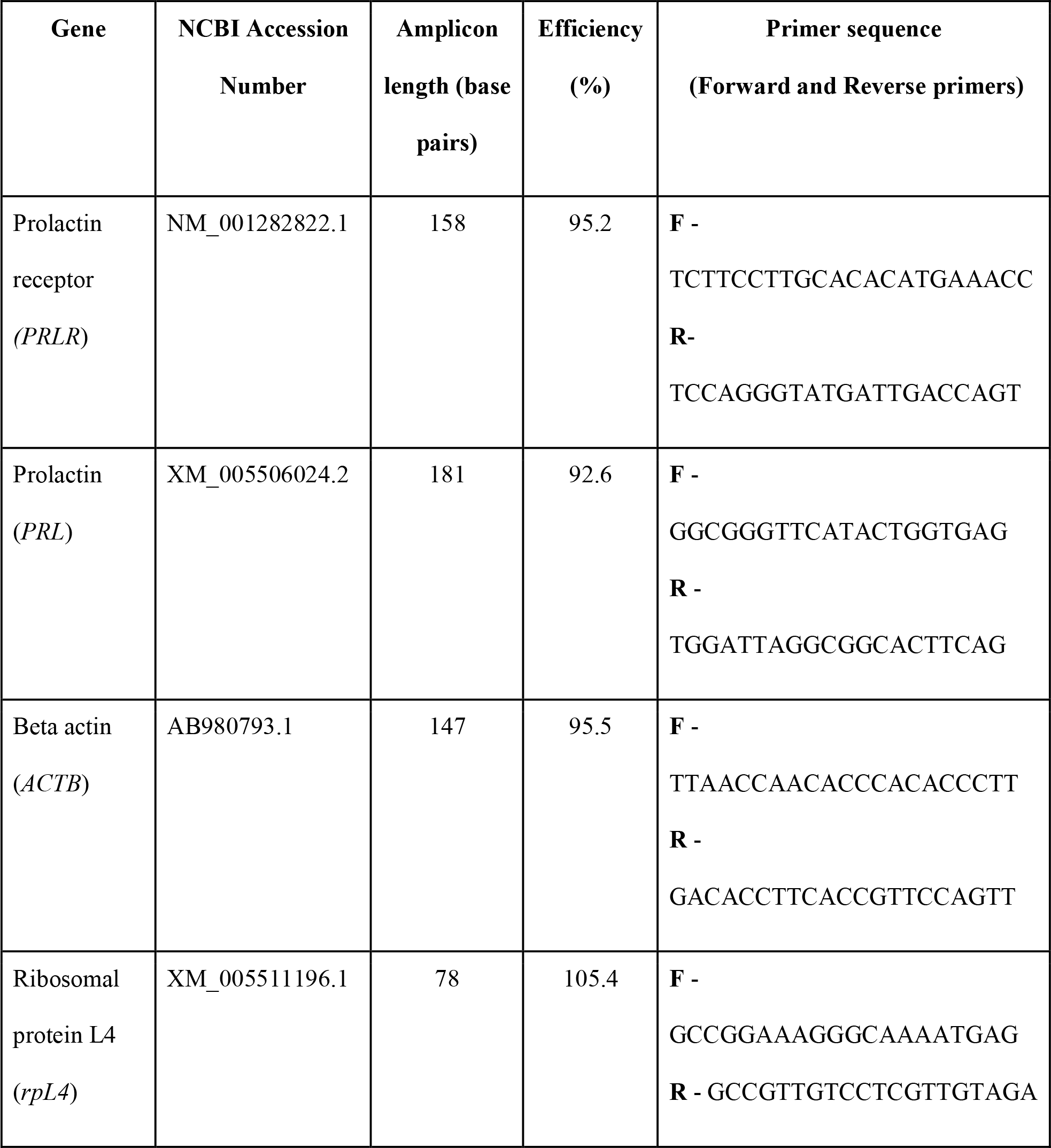
Primers used in quantitative PCR. All primers were designed using the NCBI Primer-BLAST tool using gene sequences specific to *Columba livia* (NCBI Accession numbers show the specific gene sequence from which the primers were designed). Replication efficiencies are based upon a standard curve of five 10-fold dilutions of purified gene product.

## References

1. Abs, M. (Ed.), 1983. Physiology and behaviour of the pigeon. Academic Press, London.

2. Anderson, G.M., Beijer, P., Bang, A.S., Fenwick, M.A., Bunn, S.J., Grattan, D.R., 2006. Suppression of Prolactin-Induced Signal Transducer and Activator of Transcription 5b Signaling and Induction of Suppressors of Cytokine Signaling Messenger Ribonucleic Acid in the Hypothalamic Arcuate Nucleus of the Rat during Late Pregnancy and Lactation. Endocrinology 147, 4996–5005. https://doi.org/10.1210/en.2005-0755

3. Angelier, F., Chastel, O., 2009. Stress, prolactin and parental investment in birds: A review. Gen. Comp. Endocrinol. 163, 142–148. https://doi.org/10.1016/j.ygcen.2009.03.028

4. Angelier, F., Parenteau, C., Ruault, S., Angelier, N., 2016a. Endocrine consequences of an acute stress under different thermal conditions: A study of corticosterone, prolactin, and thyroid hormones in the pigeon (Columbia livia). Comp. Biochem. Physiol. A. Mol. Integr. Physiol. 196, 38–45. https://doi.org/10.1016/j.cbpa.2016.02.010

5. Angelier, F., Weimerskirch, H., Dano, S., Chastel, O., 2007. Age, experience and reproductive performance in a long-lived bird: a hormonal perspective. Behav. Ecol. Sociobiol. 61, 611–621. https://doi.org/10.1007/s00265-006-0290-1

6. Angelier, F., Wingfield, J.C., Tartu, S., Chastel, O., 2016b. Does prolactin mediate parental and life-history decisions in response to environmental conditions in birds? A review. Horm. Behav. 77, 18–29. https://doi.org/10.1016/j.yhbeh.2015.07.014

7. Aoki, M., Wartenberg, P., Grünewald, R., Phillipps, H.R., Wyatt, A., Grattan, D.R., Boehm, U., 2019. Widespread Cell-Specific Prolactin Receptor Expression in Multiple Murine Organs. Endocrinology 160, 2587–2599. https://doi.org/10.1210/en.2019-00234

8. Austin, S., Word, K.R., 2018. Prolactin, in: Vonk, J., Shackelford, T. (Eds.), Encyclopedia of Animal Cognition and Behavior. Springer.

9. Austin, S.H., Harris, R.M., Booth, A., Lang, A.S., Farrar, V.S., Krause, J.S., Hallman, T.A., MacManes, M.D., Calisi, R.M., 2021a. Isolating the role of corticosterone in the hypothalamic-pituitary-gonadal transcriptomic stress response. Front. Endocrinol. 12.

10. Austin, S.H., Krause, J.S., Viernes, R., Farrar, V.S., Booth, A.M., Harris, R.M., Angelier, F., Lee, C., Bond, A., Wingfield, J.C., MacManes, M.D., Calisi, R.M., 2021b. Uncovering the Sex-specific Endocrine Responses to Reproduction and Parental Care. Front. Endocrinol. In press.

11. Bachelot, A., Binart, N., 2007. Reproductive role of prolactin. Reproduction 133, 361–369. https://doi.org/10.1530/REP-06-0299

12. Ball, G.F., Balthazart, J., 2008. Individual variation and the endocrine regulation of behaviour and physiology in birds: a cellular/molecular perspective. Philos. Trans. R. Soc. B Biol. Sci. 363, 1699–1710. https://doi.org/10.1098/rstb.2007.0010

13. Ben-Jonathan, N., Mershon, J.L., Allen, D.L., Steinmetz, R.W., 1996. Extrapituitary Prolactin: Distribution, Regulation, Functions, and Clinical Aspects*. Endocr. Rev. 17, 639–669. https://doi.org/10.1210/edrv-17-6-639

14. Bera, T.K., Hwang, S., Swanson, S.M., Guzman, R.C., Edery, M., Nandi, S., 1994. In situ localization of prolactin receptor message in the mammary glands of pituitaryisografted mice. Mol. Cell. Biochem. 132, 145–149. https://doi.org/10.1007/BF00926923

15. Bray, N.L., Pimentel, H., Melsted, P., Pachter, L., 2016. Near-optimal probabilistic RNA-seq quantification. Nat. Biotechnol. 34, 525–527. https://doi.org/10.1038/nbt.3519

16. Bridges, R.S., 2015. Neuroendocrine regulation of maternal behavior. Front. Neuroendocrinol. 36, 178– 196. https://doi.org/10.1016/j.yfrne.2014.11.007

17. Brown, R.S.E., Aoki, M., Ladyman, S.R., Phillipps, H.R., Wyatt, A., Boehm, U., Grattan, D.R., 2017. Prolactin action in the medial preoptic area is necessary for postpartum maternal nursing behavior. Proc. Natl. Acad. Sci. 114, 10779–10784. https://doi.org/10.1073/pnas.1708025114

18. Buntin, J., Becker, G.M., Ruzycki, E., 1991. Facilitation of parental behavior in ring doves by systemic or intracranial injections of prolactin. Horm. Behav. 25, 424–444. https://doi.org/10.1016/0018-506X(91)90012-7

19. Buntin, J.D., 1996. Neural and Hormonal Control of Parental Behavior in Birds, in: Advances in the Study of Behavior. Elsevier, pp. 161–213. https://doi.org/10.1016/S0065-3454(08)60333-2

20. Buntin, J.D., 1979. Prolactin release in parent ring doves after brief exposure to their young. J. Endocrinol. 82, 127–130. https://doi.org/10.1677/joe.0.0820127

21. Buntin, J.D., Buntin, L., 2014. Increased STAT5 signaling in the ring dove brain in response to prolactin administration and spontaneous elevations in prolactin during the breeding cycle. Gen. Comp. Endocrinol. 200, 1–9. https://doi.org/10.1016/j.ygcen.2014.02.006

22. Buntin, J.D., Ruzycki, E., 1987. Characteristics of prolactin binding sites in the brain of the ring dove (Streptopelia risoria). Gen. Comp. Endocrinol. 65, 243–253. https://doi.org/10.1016/0016-6480(87)90172-9

23. Buntin, J.D., Tesch, D., 1985. Effects of intracranial prolactin administration on maintenance of incubation readiness, ingestive behavior, and gonadal condition in ring doves. Horm. Behav. 19, 188–203. https://doi.org/10.1016/0018-506X(85)90018-2

24. Buntin, J.D., Walsh, R.J., 1988. In vivo autoradiographic analysis of prolactin binding in brain and choroid plexus of the domestic ring dove. Cell Tissue Res. 251, 105–109. https://doi.org/10.1007/BF00215453

25. Cabrera-Reyes, E.A., Vergara-Castañeda, E., Rivero-Segura, N.A., Cerbón, M., 2015. Sex differences in prolactin and its receptor expression in pituitary, hypothalamus and hippocampus of the rat. Rev. Mex. Endocrinol. Metab. Nutr. 2, 60–67.

26. Calisi, R.M., Austin, S.H., Lang, A.S., MacManes, M.D., 2018. Sex-biased transcriptomic response of the reproductive axis to stress. Horm. Behav. 100, 56–68. https://doi.org/10.1016/j.yhbeh.2017.11.011

27. Camper, P.M., Burke, W.H., 1977. The Effect of Prolactin on Reproductive Function in Female Japanese Quail (Coturnix coturnix japonica). Poult. Sci. 56, 1130–1134. https://doi.org/10.3382/ps.0561130

28. Chaiseha, Y., Ngernsoungnern, P., Sartsoongnoen, N., Prakobsaeng, N., El Halawani, M.E., 2012. Presence of prolactin mRNA in extra-pituitary brain areas in the domestic turkey. Acta Histochem. 114, 116–121. https://doi.org/10.1016/j.acthis.2011.03.007

29. Champagne, F.A., Curley, J.P., 2012. Genetics and epigenetics of parental care, in: Royle, N.J., Smiseth, P.T. (Eds.), The Evolution of Parental Care. Oxford University Press, pp. 304–324.

30. Chen, C.-C., Stairs, D.B., Boxer, R.B., Belka, G.K., Horseman, N.D., Alvarez, J.V., Chodosh, L.A., 2012. Autocrine prolactin induced by the Pten-Akt pathway is required for lactation initiation and provides a direct link between the Akt and Stat5 pathways. Genes Dev. 26, 2154–2168. https://doi.org/10.1101/gad.197343.112

31. Cheng, M., Balthazart, J., 1982. The role of nest-building activity in gonadotropin secretions and the reproductive success of ring doves (Streptopelia risoria). J. Comp. Physiol. Psychol. 96, 307–324. https://doi.org/10.1037/h0077875

32. Cheng, M.-F., Burke, W.H., 1983. Serum prolactin levels and crop-sac development in ring doves during a breeding cycle. Horm. Behav. 17, 54–65. https://doi.org/10.1016/0018-506X(83)90015-6

33. Cloues, R., Ramos, C., Silver, R., 1990. Vasoactive intestinal polypeptide-like immunoreactivity during reproduction in doves: influence of experience and number of offspring. Horm. Behav. 24, 215– 31.

34. De Vries, G.J., 2004. Minireview: Sex Differences in Adult and Developing Brains: Compensation, Compensation, Compensation. Endocrinology 145, 1063–1068. https://doi.org/10.1210/en.2003-1504

35. DeVito, W.J., 1988. Distribution of Immunoreactive Prolactin in the Male and Female Rat Brain: Effects of Hypophysectomy and Intraventricular Administration of Colchicine. Neuroendocrinology 47, 284–289. https://doi.org/10.1159/000124926

36. Dobolyi, A., Grattan, D.R., Stolzenberg, D.S., 2014. Preoptic Inputs and Mechanisms that Regulate Maternal Responsiveness. J. Neuroendocrinol. 26, 627–640. https://doi.org/10.1111/jne.12185

37. Dong, X.Y., Zhang, M., Jia, Y.X., Zou, X.T., 2012. Physiological and hormonal aspects in female domestic pigeons (*Columba livia*) associated with breeding stage and experience: Pigeon physiological and hormonal changes. J. Anim. Physiol. Anim. Nutr. no-no. https://doi.org/10.1111/j.1439-0396.2012.01331.x

38. El Halawani, M.E., Silsby, J.L., Behnke, E.J., Fehrer, S.C., 1986. Hormonal Induction of Incubation Behavior in Ovariectomized Female Turkeys (Meleagris Gallopavo). Biol. Reprod. 35, 59–67. https://doi.org/10.1095/biolreprod35.1.59

39. Featherstone, K., White, M.R.H., Davis, J.R.E., 2012. The Prolactin Gene: A Paradigm of Tissue Specific Gene Regulation with Complex Temporal Transcription Dynamics. J. Neuroendocrinol. 24, 977–990. https://doi.org/10.1111/j.1365-2826.2012.02310.x

40. Feder, H.H., Storey, A., Goodwin, D., Reboulleau, C., Silver, R., 1977. Testosterone and “5 -Dihydrotestosterone” Levels in Peripheral Plasma of Male and Female Ring Doves (Streptopelia risoria) During the Reproductive Cycle1. Biol. Reprod. 16, 666–677. https://doi.org/10.1095/biolreprod16.5.666

41. Ferraris, J., Boutillon, F., Bernadet, M., Seilicovich, A., Goffin, V., Pisera, D., 2012. Prolactin receptor antagonism in mouse anterior pituitary: effects on cell turnover and prolactin receptor expression. Am. J. Physiol.-Endocrinol. Metab. 302, E356–E364. https://doi.org/10.1152/ajpendo.00333.2011

42. Freeman, M.E., Kanyicska, B., Lerant, A., Nagy, G., 2000. Prolactin: Structure, Function, and Regulation of Secretion. Physiol. Rev. 80, 1523–1631. https://doi.org/10.1152/physrev.2000.80.4.1523

43. Gillespie, M.J., Crowley, T.M., Haring, V.R., Wilson, S.L., Harper, J.A., Payne, J.S., Green, D., Monaghan, P., Donald, J.A., Nicholas, K.R., Moore, R.J., 2013. Transcriptome analysis of pigeon milk production – role of cornification and triglyceride synthesis genes. BMC Genomics 14, 169. https://doi.org/10.1186/1471-2164-14-169

44. Gill-Sharma, M.K., 2009. Prolactin and Male Fertility: The Long and Short Feedback Regulation. Int. J. Endocrinol. 2009, 1–13. https://doi.org/10.1155/2009/687259

45. Grattan, D.R., 2018. Coordination or Coincidence? The Relationship between Prolactin and Gonadotropin Secretion. Trends Endocrinol. Metab. 29, 3–5. https://doi.org/10.1016/j.tem.2017.11.004

46. Grattan, D.R., Jasoni, C.L., Liu, X., Anderson, G.M., Herbison, A.E., 2007. Prolactin Regulation of Gonadotropin-Releasing Hormone Neurons to Suppress Luteinizing Hormone Secretion in Mice. Endocrinology 148, 4344–4351. https://doi.org/10.1210/en.2007-0403

47. Grattan, D.R., Le Tissier, P., 2015. Hypothalamic control of prolactin secretion, and the multiple reproductive functions of prolactin, in: Plant, T.M., Zeleznik, A.J., Knobil, E., Neil, J.D. (Eds.), Knobil and Neill’s Physiology of Reproduction. Elsevier/Academic Press, Amsterdam, pp. 469– 526.

48. Guillaumot, P., Benahmed, M., 1999. Prolactin receptors are expressed and hormonally regulated in rat Sertoli cells. Mol. Cell. Endocrinol. 149, 163–168. https://doi.org/10.1016/S0303-7207(98)00246-9

49. Hansen, E.W., 1966. Squab-induced crop growth in ring dove foster parents. J. Comp. Physiol. Psychol. 62, 120–122. https://doi.org/10.1037/h0023477

50. Hennighausen, L., Robinson, G.W., Wagner, K.-U., Liu, X., 1997. Prolactin Signaling in Mammary Gland Development. J. Biol. Chem. 272, 7567–7569. https://doi.org/10.1074/jbc.272.12.7567

51. Hope, S.F., DuRant, S.E., Angelier, F., Hallagan, J.J., Moore, I.T., Parenteau, C., Kennamer, R.A., Hopkins, W.A., 2020. Prolactin is related to incubation constancy and egg temperature following a disturbance in a precocial bird. Gen. Comp. Endocrinol. 295, 113489. https://doi.org/10.1016/j.ygcen.2020.113489

52. Horseman, N.D., Buntin, J.D., 1995. Regulation of Pigeon Cropmilk Secretion and Parental Behaviors by Prolactin. Annu. Rev. Nutr. 15, 213–238. https://doi.org/10.1146/annurev.nu.15.070195.001241

53. Hu, S., Duggavathi, R., Zadworny, D., 2017. Regulatory Mechanisms Underlying the Expression of Prolactin Receptor in Chicken Granulosa Cells. PLOS ONE 12, e0170409. https://doi.org/10.1371/journal.pone.0170409

54. Hutchison, R.E., 1975. EFFECTS OF OVARIAN STEROIDS AND PROLACTIN ON THE SEQUENTIAL DEVELOPMENT OF NESTING BEHAVIOUR IN FEMALE BUDGERIGARS. J. Endocrinol. 67, 29–39. https://doi.org/10.1677/joe.0.0670029

55. Ketterson, E.D., Atwell, J.W., McGlothlin, J.W., 2009. Phenotypic integration and independence: Hormones, performance, and response to environmental change. Integr. Comp. Biol. 49, 365–379. https://doi.org/10.1093/icb/icp057

56. Kurima, K., Proudman, J.A., El Halawani, M.E., Wong, E.A., 1995. The turkey prolactin-encoding gene and its regulatory region. Gene 156, 309–310. https://doi.org/10.1016/0378-1119(95)00032-2

57. Lea, R.W., Vowles, D.M., 1985. The control of prolactin secretion and nest defence in the ring dove (*Streptopelia risoria)*. Bolletino Zool. 52, 323–329. https://doi.org/10.1080/11250008509440534

58. Lea, R.W., Vowles, D.M., Dick, H.R., 1986. Factors affecting prolactin secretion during the breeding cycle of the ring dove (Streptopelia risoria) and its possible role in incubation. J. Endocrinol. 110, 447–458. https://doi.org/10.1677/joe.0.1100447

59. Livak, K.J., Schmittgen, T.D., 2001. Analysis of Relative Gene Expression Data Using Real-Time Quantitative PCR and the 2−ΔΔCT Method. Methods 25, 402–408. https://doi.org/10.1006/meth.2001.1262

60. Lopez Vicchi, F., Ladyman, S.R., Ornstein, A.M., Gustafson, P., Knowles, P., Luque, G.M., Grattan, D.R., Becu Villalobos, D., 2020. Chronic high prolactin levels impact on gene expression at discrete hypothalamic nuclei involved in food intake. FASEB J. 34, 3902–3914. https://doi.org/10.1096/fj.201902357R

61. Love, M.I., Huber, W., Anders, S., 2014. Moderated estimation of fold change and dispersion for RNA-seq data with DESeq2. Genome Biol. 15, 550. https://doi.org/10.1186/s13059-014-0550-8

62. MacManes, M.D., Austin, S.H., Lang, A.S., Booth, A., Farrar, V., Calisi, R.M., 2017. Widespread patterns of sexually dimorphic gene expression in an avian hypothalamic–pituitary–gonadal (HPG) axis. Sci. Rep. 7, 45125. https://doi.org/10.1038/srep45125

63. Macnamee, M.C., Sharp, P.J., Lea, R.W., Sterling, R.J., Harvey, S., 1986. Evidence that vasoactive intestinal polypeptide is a physiological prolactin-releasing factor in the bantam hen. Gen. Comp. Endocrinol. 62, 470–478. https://doi.org/10.1016/0016-6480(86)90057-2

64. Marano, R.J., Ben-Jonathan, N., 2014. Minireview: Extrapituitary Prolactin: An Update on the Distribution, Regulation, and Functions. Mol. Endocrinol. 28, 622–633. https://doi.org/10.1210/me.2013-1349

65. Maurer, R.A., Notides, A.C., 1987. Identification of an estrogen-responsive element from the 5’-flanking region of the rat prolactin gene. Mol. Cell. Biol. 7, 4247–4254. https://doi.org/10.1128/MCB.7.12.4247

66. McCarthy, M.M., Konkle, A.T.M., 2005. When is a sex difference not a sex difference? Front. Neuroendocrinol. 26, 85–102. https://doi.org/10.1016/j.yfrne.2005.06.001

67. Meier, A.H., 1969. Antigonadal effects of prolactin in the White-throated Sparrow (Zonotrichia albicollis). Gen. Comp. Endocrinol. 13, 222–225.

68. Meier, A.H., Martin, D.D., MacGregor, R., 1971. Temporal synergism of corticosterone and prolactin controlling gonadal growth in sparrows. Science 173, 1240–1242.

69. Michel, G.F., 1977. Experience and progesterone in ring dove incubation. Anim. Behav. 25, 281–285. https://doi.org/10.1016/0003-3472(77)90003-3

70. Moult, P.J., Besser, G., 1981. Prolactin and gonad function. IPPF Med Bull 15, 3–4.

71. Nagano, M., Kelly, P., 1999. Tissue distribution and regulation of rat prolactin receptor gene expression. Quantitative analysis by polymerase chain reaction. J. Biol. Chem. 269, 13337–45.

72. Pi, X., Voogt, J.L., 2002. Sex difference and estrous cycle: expression of prolactin receptor mRNA in rat brain. Mol. Brain Res. 103, 130–139. https://doi.org/10.1016/S0169-328X(02)00194-8

73. Pitts, G.R., Youngren, O.M., Silsby, J.L., Rozenboim, I., Chaiseha, Y., Phillips, R.E., Foster, D.N., El Halawani, M.E., 1994. Role of Vasoactive Intestinal Peptide in the Control of Prolactin-Induced Turkey Incubation Behavior. II. Chronic Infusion of Vasoactive Intestinal Peptide1. Biol. Reprod. 50, 1350–1356. https://doi.org/10.1095/biolreprod50.6.1350

74. Ramesh, R., Kuenzel, W.J., Buntin, J.D., Proudman, J.A., 2000. Identification of growth-hormone and prolactin-containing neurons within the avian brain. Cell Tissue Res. 299, 317–383. https://doi.org/10.1007/s004419900104

75. Ramsey, S.M., Goldsmith, A.R., Silver, R., 1985. Stimulus requirements for prolactin and LH secretion in incubating ring doves. Gen. Comp. Endocrinol. 59, 246–256. https://doi.org/10.1016/0016-6480(85)90376-4

76. Reddy, I.J., David, C.G., Sarma, P.V., Singh, K., 2002. The possible role of prolactin in laying performance and steroid hormone secretion in domestic hen (Gallus domesticus). Gen. Comp. Endocrinol. 127, 249–255. https://doi.org/10.1016/S0016-6480(02)00034-5

77. Riddle, O., Braucher, P.F., 1931. Studies on the Physiology of Reproduction in Birds: XXX. Control of the Special Secretion of the Crop-Gland in Pigeons by an Anterior Pituitary Hormone. Am. J. Physiol.-Leg. Content 97, 617–625. https://doi.org/10.1152/ajplegacy.1931.97.4.617

78. Rozenboim, I., Tabibzadeh, C., Silsby, J.L., El Halawani, M.E., 1993. Effect of Ovine Prolactin Administration on Hypothalamic Vasoactive Intestinal Peptide (VIP), Gonadotropin Releasing Hormone I and II Content, and Anterior Pituitary VIP Receptors in Laying Turkey Hens1. Biol. Reprod. 48, 1246–1250. https://doi.org/10.1095/biolreprod48.6.1246

79. Salais-López, H., Agustín-Pavón, C., Lanuza, E., Martínez-García, F., 2018. The maternal hormone in the male brain: Sexually dimorphic distribution of prolactin signalling in the mouse brain. PLOS ONE 13, e0208960. https://doi.org/10.1371/journal.pone.0208960

80. Shani, J., Barkey, R.J., Amit, T., 1981. Endogenous Prolactin Maintains its own Binding Sites in the Pigeon Crop Sac Mucosa. J. Recept. Res. 2, 407–417. https://doi.org/10.3109/107998981809038875

81. Silver, R., 1978. The Parental Behavior of Ring Doves: The intricately coordinated behavior of the male and female is based on distinct physiological mechanisms in the sexes. Am. Sci. 66, 209–215.

82. Sjoeholm, A., Bridges, R.S., Grattan, D.R., Anderson, G.M., 2011. Region-, Neuron-, and Signaling Pathway-Specific Increases in Prolactin Responsiveness in Reproductively Experienced Female Rats. Endocrinology 152, 1979–1988. https://doi.org/10.1210/en.2010-1220

83. Slawski, B.A., Buntin, J.D., 1995. Preoptic area lesions disrupt prolactin-induced parental feeding behavior in ring doves. Horm. Behav. 29, 248–266.

84. Smiley, K.O., 2019. Prolactin and avian parental care: New insights and unanswered questions. Horm. Behav. https://doi.org/10.1016/j.yhbeh.2019.02.012

85. Smiley, K.O., Buntin, J.D., Corbitt, C., Deviche, P., 2020. Central prolactin binding site densities change seasonally in an adult male passerine bird (Junco hyemalis). J. Chem. Neuroanat. 106, 101786. https://doi.org/10.1016/j.jchemneu.2020.101786

86. Smiley, K.O., Dong, L., Ramakrishnan, S., Adkins-Regan, E., 2021. Central prolactin receptor distribution and pSTAT5 activation patterns in breeding and non-breeding zebra finches (Taeniopygia guttata). Gen. Comp. Endocrinol. 301, 113657. https://doi.org/10.1016/j.ygcen.2020.113657

87. Sockman, K.W., Schwabl, H., Sharp, P.J., 2000. The Role of Prolactin in the Regulation of Clutch Size and Onset of Incubation Behavior in the American Kestrel. Horm. Behav. 38, 168–176. https://doi.org/10.1006/hbeh.2000.1616

88. Soneson, C., Love, M.I., Robinson, M.D., 2016. Differential analyses for RNA-seq: transcript-level estimates improve gene-level inferences. F1000Research 4, 1521. https://doi.org/10.12688/f1000research.7563.2

89. Stiver, K.A., Alonzo, S.H., 2009. Parental and Mating Effort: Is There Necessarily a Trade-Off? Ethology 115, 1101–1126. https://doi.org/10.1111/j.1439-0310.2009.01707.x

90. Swaminathan, G., Varghese, B., Fuchs, S.Y., 2008. Regulation of Prolactin Receptor Levels and Activity in Breast Cancer. J. Mammary Gland Biol. Neoplasia 13, 81–91. https://doi.org/10.1007/s10911-008-9068-6

91. Torner, L., Maloumby, R., Nava, G., Aranda, J., Clapp, C., Neumann, I.D., 2004. In vivo release and gene upregulation of brain prolactin in response to physiological stimuli. Eur. J. Neurosci. 19, 1601– 1608. https://doi.org/10.1111/j.1460-9568.2004.03264.x

92. Torner, L., Toschi, N., Nava, G., Clapp, C., Neumann, I.D., 2002. Increased hypothalamic expression of prolactin in lactation: involvement in behavioural and neuroendocrine stress responses: Prolactin modulates stress responses during lactation. Eur. J. Neurosci. 15, 1381–1389. https://doi.org/10.1046/j.1460-9568.2002.01965.x

93. Vleck, C.M., Patrick, D.J., 1999. Effects of Vasoactive Intestinal Peptide on Prolactin Secretion in Three Species of Passerine Birds. Gen. Comp. Endocrinol. 113, 146–154. https://doi.org/10.1006/gcen.1998.7191

94. Zera, A.J., Harshman, L.G., 2001. The Physiology of Life History Trade-Offs in Animals. Annu. Rev. Ecol. Syst. 32, 95–126. https://doi.org/10.1146/annurev.ecolsys.32.081501.114006

95. Zhou, J.F., Zadworny, D., Gueméné, D., Kuhnlein, U., 1996. Molecular Cloning, Tissue Distribution, and Expression of the Prolactin Receptor during Various Reproductive States in Meleagris gallopavo 1. Biol. Reprod. 55, 1081–1090. https://doi.org/10.1095/biolreprod55.5.1081

96. Zinzow-Kramer, W.M., Horton, B.M., Maney, D.L., 2014. Evaluation of reference genes for quantitative real-time PCR in the brain, pituitary, and gonads of songbirds. Horm. Behav. 66, 267–275. https://doi.org/10.1016/j.yhbeh.2014.04.011

