## Supplemental Table 1 for "Prolactin and prolactin receptor expression in the HPG axis and crop during parental care in both sexes of a biparental bird (*Columba livia*)"

| **Stage** | **Sex** | **HPG RNAseq data (n)** | **Crop (total) (n)** | **Crops without associated RNAseq data (n)** |
| --- | --- | --- | --- | --- |
| Nest building | F | 10 | 10 | 4 |
|  | M | 10 | 8 | 4 |
| Clutch completion  (incubation day 3) | F | 10 | 10 | 1 |
|  | M | 10 | 9 | 0 |
| Mid-incubation  (incubation day 9) | F | 12 | 9 | 0 |
|  | M | 10 | 10 | 2 |
| Early Hatch manipulation (manipulation on incubation day 8) | F | 10 | 8 | 0 |
|  | M | 10 | 8 | 0 |
| Hatching | F | 10 | 9 | 4 |
|  | M | 10 | 11 | 5 |
