## Supplemental Table 2 for "Prolactin and prolactin receptor expression in the HPG axis and crop during parental care in both sexes of a biparental bird (*Columba livia*)"

| **Gene** | **NCBI Accession Number** | **Amplicon length (base pairs)** | **Efficiency (%)** | **Primer sequence**  **(Forward and Reverse primers)** |
| --- | --- | --- | --- | --- |
| Prolactin receptor *(PRLR*) | NM_001282822.1 | 158 | 95.2 | **F** - TCTTCCTTGCACACATGAAACC  **R**- TCCAGGGTATGATTGACCAGT |
| Prolactin (*PRL*) | XM_005506024.2 | 181 | 92.6 | **F** - GGCGGGTTCATACTGGTGAG  **R** - TGGATTAGGCGGCACTTCAG |
| Beta actin (*ACTB*) | AB980793.1 | 147 | 95.5 | **F** - TTAACCAACACCCACACCCTT  **R** - GACACCTTCACCGTTCCAGTT |
| Ribosomal protein L4 (*rpL4*) | XM_005511196.1 | 78 | 105.4 | **F** - GCCGGAAAGGGCAAAATGAG  **R** - GCCGTTGTCCTCGTTGTAGA |
